# Shared striatal activity in decisions to satisfy curiosity and hunger at the risk of electric shocks

**DOI:** 10.1101/473975

**Authors:** Johnny King L Lau, Hiroki Ozono, Kei Kuratomi, Asuka Komiya, Kou Murayama

## Abstract

Curiosity is often portrayed as a desirable feature of human faculty. However, curiosity may come at a cost that sometimes puts people in a harmful situation. Here, with a set of behavioural and neuroimaging experiments using stimuli that strongly trigger curiosity (e.g., magic tricks), we examined the psychological and neural mechanisms underlying the motivational effect of curiosity. We consistently demonstrated that across different samples, people were indeed willing to gamble, subjecting themselves to physical risks (i.e. electric shocks) in order to satisfy their curiosity for trivial knowledge that carries no apparent instrumental value. Also, this influence of curiosity shares common neural mechanisms with that of extrinsic incentives (i.e. hunger for food). In particular, we showed that acceptance (compared to rejection) of curiosity/incentive-driven gambles was accompanied by enhanced activity in the ventral striatum (when curiosity was elicited), which extended into the dorsal striatum (when participants made a decision).

## Introduction

Curiosity is a fundamental part of human motivation that supports an enormous variety of human intellectual behaviours, ranging from early learning in children to scientific discovery ^1–3^. A meta-analysis indicates that intellectual curiosity predicts academic performance over and above intelligence ^4^, and corroborating findings show the benefits of curiosity in enhancing long-term consolidation of learning and memory ^5,6^. The critical importance of curiosity in human intellectual behaviour is succinctly expressed in Albert Einstein’s famous quote “I have no special talent. I am only passionately curious.” Moreover, empirical literature has suggested a number of positive outcomes associated with curiosity over the life span ^4,7–9^. However, in both historic and modern literature, the positive portrayal of curiosity is often compromised by its inherent negative aspect: strong seductive nature ^10^. In Greek mythology, for example, after losing his beloved Eurydice to the underworld, Orpheus convinced the gods to let him take her back to the world of the living on the condition that he would not look back until they had returned. Orpheus could not help but look back; he succumbed to curiosity and lost Eurydice. This theme, which appears repeatedly in classic ancient anecdotes (e.g., Pandora, Psyche, Eve), illustrates the motivational power of curiosity, which biases our decision making despite the knowledge of consequential negative outcomes.

How does curiosity for inconsequential knowledge motivate people to make risky decisions? Although there have been various theories on the construct and origin of curiosity in the literature ^10–12^, a recent body of research has seen an emerging consensus that, like food and other extrinsic rewards, curiosity can be understood as a reward-learning process of knowledge acquisition or information seeking ^8,13–16^. According to this account, individuals are motivated to actively seek knowledge because knowledge acquisition serves as an inherent ‘reward’ (possibly due to uncertainty reduction and potential anticipatory utility^17^), reinforcing further information seeking or behaviours that promote information gain. In fact, studies have repeatedly demonstrated that both animals ^18–20^ and humans ^6,21–23^ are willing to pay small amounts to satisfy their curiosity for knowledge about a future reward that cannot be changed. For example, people are willing to sacrifice parts of future monetary reward in exchange for immediate information just to learn about the outcome of a monetary lottery, even though that information cannot be used to alter prospective lottery outcomes^21^.

Past work on how extrinsic incentives shape behaviour and decisions highlight the involvement of a strong, visceral motivational feeling, known as incentive salience (‘wanting’) ^24,25^. Incentive salience can readily be triggered by encountering reward cues and by vivid imagery of reward. Also, it is hypothesised to integrate the current physiological neurobiological state and previously learned associations about the reward. For example, when you arrive hungry to a restaurant, you may start salivating and find every item on the menu to be more appealing than they usually are to you. The motivational urge of incentive salience may even prompt people towards irrational, impulsive and even obsessive behaviours^26,27^. Previous animal research suggested that incentive salience is mediated by the mesolimbic dopamine system as well as the surrounding relatively large brain network including both cortical and subcortical areas ^28,29^, whereas neuroimaging studies in human have identified the particular involvement of the ventral striatum (i.e. the nucleus accumbens) and the dorsal striatum (i.e. the caudate) in the processing of different extrinsic rewards ^30–32^. For example, Lawrence et al. showed that the activation in the nucleus accumbens in response to food cues predicts participants’ consumption of crisps when they are left alone in a room after the experiment ^33^.

Integrating these ideas and findings, we suggest that, like extrinsic incentives such as food, the rewarding property of knowledge may evoke incentive salience, energising people toward knowledge acquisition behaviour even if it entails significant risks ^15,34^. In fact, although the neurobiological mechanisms underpinning curiosity-based decisions have been under-examined in the literature, the investigations using brain imaging collectively suggest that the states of curiosity about information that does not signal any future rewards modulate neural regions in dopamine systems, including caudate nucleus, nucleus accumbens and midbrain structures (i.e. ventral tegmental area/substantia nigra [VTA/SN])^5,6,35^. As discussed previously, these areas, with the caudate nucleus and nucleus accumbens in particular, have been implicated in incentive salience in neuroimaging studies of cognitive processing related to food stimuli^30^. A recent study, using a neurocomputational approach and a monetary gamble paradigm, demonstrated that the same patterns of activations in the striatum responding to the values of potential rewards (i.e. the varying amount of monetary gain) can also predict subjective value of the information (i.e. how much a person would be willing to pay for extra information), which may be partially driven by non-instrumental, curiosity-based motives^36^. Taking these together, it is possible that decisions driven by a strong urge of knowledge acquisition rely on similar mechanisms that support motivated behaviours triggered by incentive salience of extrinsic incentives^37,38^.

The current study examines whether and how people are willing to subject themselves to physical risks to satisfy their curiosity for trivial, inconsequential knowledge, by using a ‘curiosity-based risky decision-making’ paradigm in a series of behavioural and fMRI experiments. In the ‘curiosity condition’, participants first saw a curiosity-stimulating stimulus (i.e. video clips of magic tricks, or text-based trivia questions), followed by a wheel of fortune which visually depicted the probability of winning (versus losing) a lottery in each trial. Participants then had to make a decision regarding whether or not they would gamble on the lottery (risking electric shock) to have a chance of learning the solution to the trick. The experiment also included food trials, in which participants were presented with food pictures and decided whether to gamble (risking electric shock) to have a chance of obtaining the food. The inclusion of the food condition allows us to directly compare the influence of curiosity and extrinsic incentives on risk-based decisions and to examine the common and differentiated mechanisms supporting them. We postulated that i) people are willing to subject themselves to physical risks (i.e. electric shocks) to satisfy their curiosity for trivial, inconsequential knowledge (i.e. magic tricks, trivia) or their hunger for food, and ii) the motivational lure of curiosity and hunger share common neural correlates implicated in the incentive salience of food, namely the ventral and dorsal striatum ^30^.

## Results

In the decision-making task, participants viewed short video clips of magic tricks (curiosity condition) performed by professional magicians (filmed specifically for this study; see Supplementary Videos 1-2), followed by a wheel of fortune visually depicting the outcome probability of winning (versus losing) a lottery (with a certain probability; between 16.7% - 83.3%; see Methods) in each trial (Fig. 1). Participants were then asked to make a decision regarding whether they would gamble on the lottery or not. They were told that, if they accepted to gamble and won, they might see the secret behind the magic trick after the experiment as a reward. The experiment also included a food condition, in which participants were presented with food pictures and were told that they might have the food as a reward after the experiment if they accepted to gamble and won. Critically, for both types of trials, if participants gambled and lost, they would expect to receive electric shock after the experiment. The magnitude of electric shock was calibrated and demonstrated to participants before the experimental task. Stimulating electrodes were attached to the participants throughout the task and the shock was expected to be delivered at the very end. For each stimulus, participants gave a rating of how curious they were to see the solution to the trick (curiosity rating) or how much they would like to eat the food item (desirability rating). To maximise participants’ desire for food during the experiment, they were required to refrain from food consumption shortly before attending the testing session. Additional details on the task are reported in the Methods section: see ‘Procedure’.

**Fig. 1:**
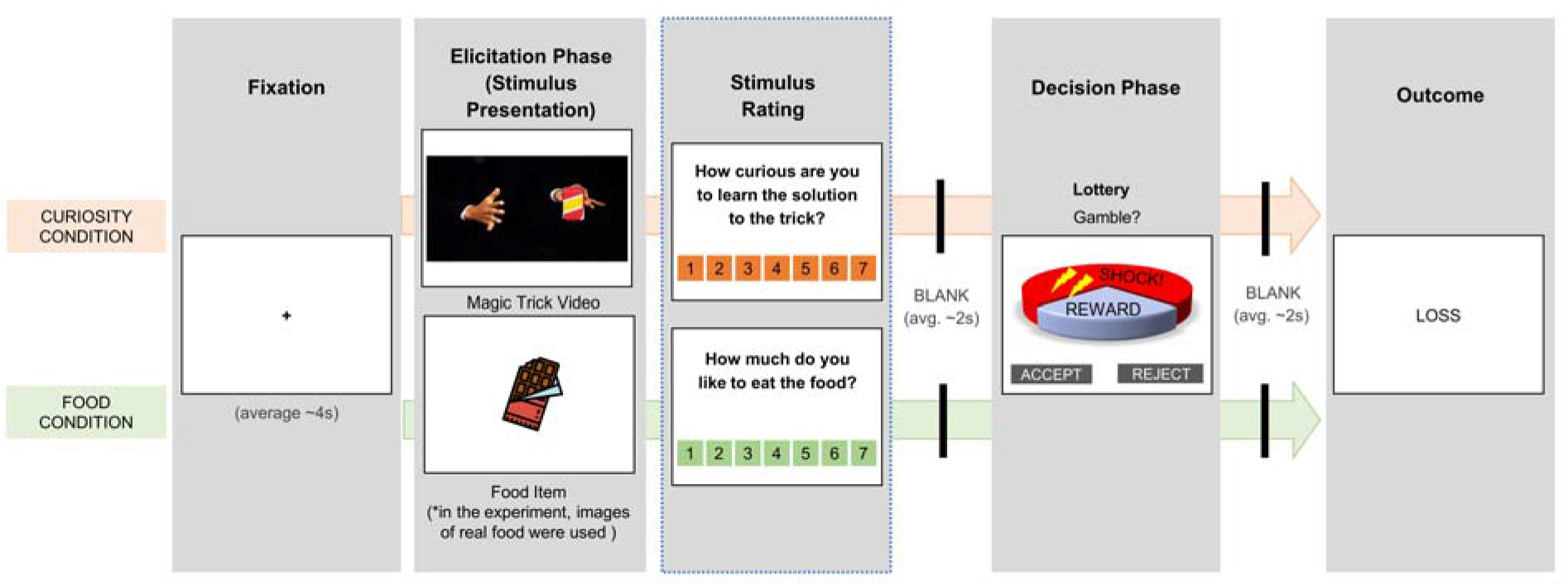
Experimental task. In each experimental trial, participants viewed either a video of a magic trick (curiosity condition) or an image of a food item (food condition). They then gave a rating to indicate their level of curiosity about the magic trick or the level of desirability of the food. After rating the stimulus, participants were shown a wheel of fortune (WoF) representing a lottery, which visualised the probability of them winning/losing in that trial (in this example, as visually presented in the WoF, there is a higher probability of winning compared with losing). Participants had to decide whether to gamble or not. Participants were instructed that if they accepted the gamble and won, they might see the secret behind the magic trick (or get the food, for food condition) after the experiment as a reward. If they gambled and lost, they would expect to receive electric shock after the experiment. Participants could also opt to reject the gamble. Participants understood that, in general, the more gambles they won, the more likely they were to see more solutions to the tricks and obtain more food items. Similarly, the more gambles they lost, the more likely they were to receive electric shock for a longer duration. In the modified version of the task for fMRI experiments, (due to time constraints in scanning) the rating phase appeared in only 10% of all the trials during the task. In addition, for the ‘trivia version’ of the fMRI experiment, participants were presented with trivia questions (instead of magic tricks) as stimuli in the curiosity condition.

### Behavioural experiments

#### Curiosity and food desirability bias decision-making

In the initial behavioural experiment (N = 32), generalised linear mixed-effects modelling (GLME) on all trials showed that participants were more likely to reject the gamble as the presented outcome probability of receiving electric shock increased [GLME: *Z* = −19.045, *P* < 0.001, Exp(β) = 0.455, β = −0.787, 95% C.I.◻=◻−0.868 – −0.706]. This indicates that electric shock worked effectively as an aversive stimulus. Importantly, above and beyond the shock probability, stimulus rating (i.e. curiosity or food desirability rating) positively predicted the ‘accept’ decision [GLME: *Z* = 20.253, *P* < 0.001, Exp (β) = 2.110, β =0.747, 95% C.I. = 0.674 – 0.819]. The main effect of ‘Incentive Category’ (curiosity vs. food) on the decision did not attain the set threshold of statistical significance [GLME: *Z* = 1.25, *P* = 0.211, Exp(β) = 1.067, β = 0.065, 95% C.I.◻=◻−0.037 – 0.167]. Also, none of the interaction effects between the stimulus rating, presented probability, and incentive category (including a three-way interaction) reached the set threshold of statistical significance (all *P*s > 0.05). Thus, there is no evidence the observed association between rating and acceptance rate differ between different categories and levels of shock probability. In fact, when we analysed curiosity trials and food trials separately, in addition to outcome probability, ratings of curiosity and food desirability were similarly associated with higher tendency to accept the gamble [GLME - curiosity: Z=7.236, Exp(β) = 3.185, β = 1.159, 95% C.I. = 0.845 – 1.472; food: Z=7.493, Exp(β) = 3.145, β = 1.146, 95% C.I. = 0.846 – 1.446; *Ps* < 0.001 in both conditions] (Fig. 2; also see Supplementary Table 1).

**Fig. 2:**
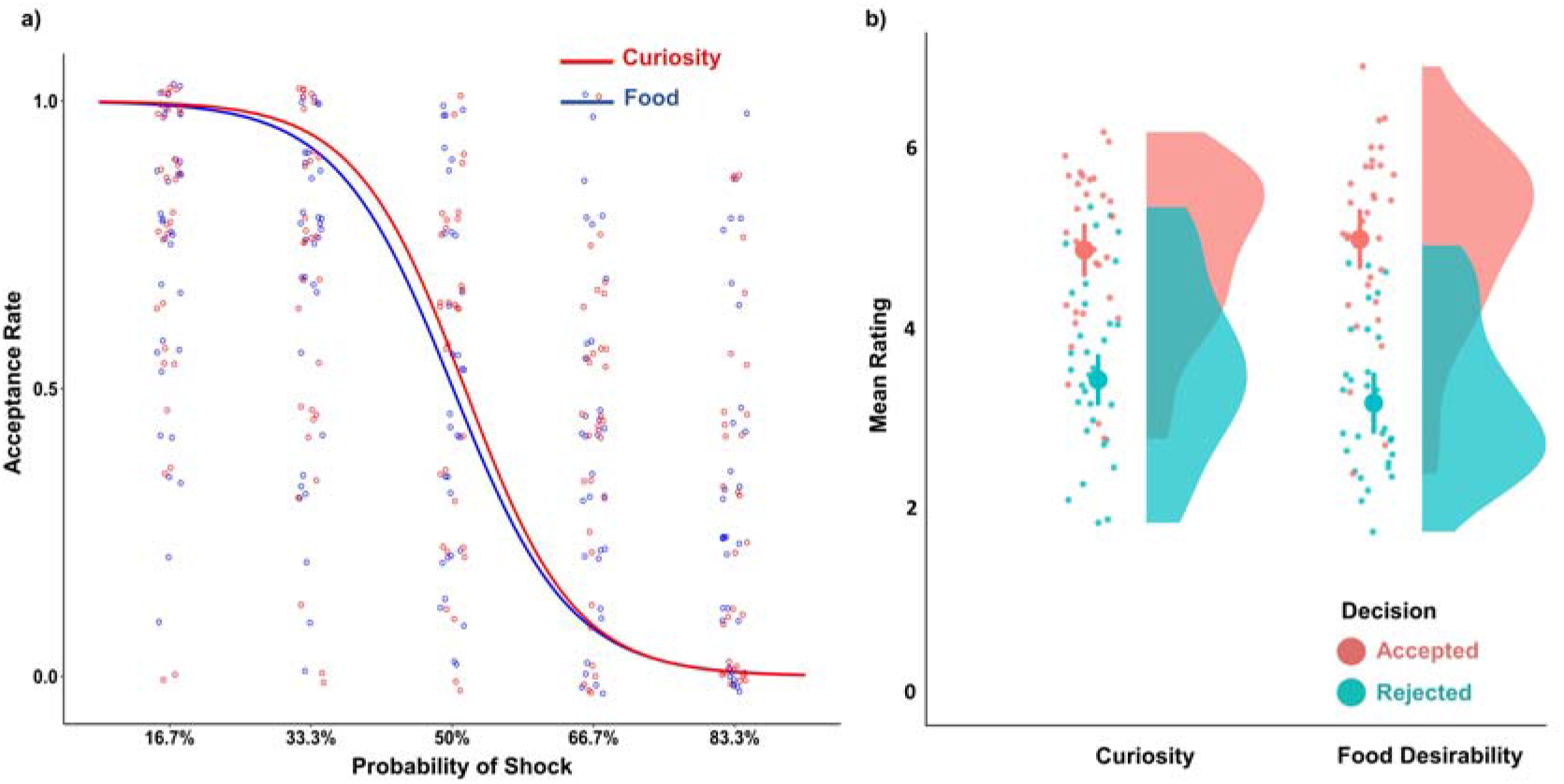
Behavioural results of motivation-driven decision-making. Both graphs include data of 32 participants from the initial behavioural experiment. a) As illustrated by the modelled logistic curve, participants tended to reject the gamble more as the probability of electric shock presented in a wheel of fortune increased, in curiosity (Z=−7.80, P<0.001, β=−1.28; red curve) and food (Z=−6.29, P<0.001, β=−1.21; blue curve) conditions. Acceptance rate on y-axis ranges from 0 (100% rejection) to 1 (100% acceptance). Each dot represents the average acceptance rate of a participant at that level of shock probability presented in each condition. b) As illustrated in the raincloud plot on the right, higher average ratings of curiosity about magic tricks (Z=7.24, P<0.001, β=1.16) and food desirability (Z=7.49, P<0.001, β=1.15) were shown for accepted (orange solid circle) compared with rejected (blue solid circle) trials. Each dot represents the average rating per participant, for each condition. The split-half violin plots show the probability density of the data at different ratings. Error bars indicate standard error of the mean. In both graphs, dots are jittered for better visual display. Statistical significance was assessed with generalised linear mixed-effects models (see ‘Behavioural analysis’ in Methods).

#### Did curiosity distort the perception of the probability of outcome?

As shown in the initial experiment, people are willing to subject themselves to potential risks to satisfy their curiosity for trivial, inconsequential knowledge, corroborating our hypothesis. However, an alternative explanation may be that curiosity simply distorts one’s perception of the probability of risks. People tend to report higher probabilities for positive events and lower probabilities for negative events when they are in a pleasant mood ^39^. It could be that when people are curious, feel more motivated and experience positive feelings, they overestimate the probability of winning in the lottery (or underestimate the risk). To evaluate this possibility, we carried out a follow-up behavioural experiment. In this experiment, after seeing a curiosity-/hunger-evoking stimulus, participants (N = 29) estimated their subjective chance of winning(/losing) in the lottery when presented with a wheel of fortune in each trial, before also making their actual decision to gamble or not (see Supplementary Methods for the procedures of this modified task).

First, we applied the same GLME to predict participants’ decision as an attempt to check whether the results from the initial experiment could be replicated. In this follow-up experiment, in addition to the presented shock probability [GLME: *Z* = −17.334, *P* < 0.001, Exp(β) = 0.251, β = −1.381, 95% C.I.◻=◻−1.537 – −1.225], stimulus rating [GLME: *Z* = 19.568, *P* < 0.001, Exp (β) = 3.947, β = 1.373, 95% C.I. = 1.236 – 1.511] again positively predicted the decision to gamble. There was no statistically significant main effect of ‘Incentive Category’ (curiosity vs. food) on the decision [GLME: *Z* = −0.505, *P* = 0.613, Exp (β) = 0.958, β = −0.043, 95% C.I. = −0.208 − 0.123]. Also, none of the interaction effects between the presented probability, stimulus rating, and incentive category (including a three-way interaction) reached the set threshold of statistical significance (all *P*s > 0.05). These results are consistent with those from the initial experiment (see Supplementary Table 1, for separate analysis of curiosity and food conditions).

As the main objective of the follow-up experiment, we analysed the predictors of participants' subjective estimation of the chance to win/lose with a linear mixed-effects model (LME). Participant’s own estimation of win/loss probability was predicted by the presented shock (outcome) probability that they actually viewed in each trial [LME - with all trials: t = − 27.522, β = −1.762, 95% C.I. = −1.887 – −1.637; with curiosity trials only: t = −25.519, β = −1.798, 95% C.I. = −1.936 – −1.660; with food trials only: t = −26.27, β = −1.728, 95% C.I. = −1.857 – −1.599; *Ps* < 0.001 in all cases], which perhaps is not too surprising. In contrast, there was no evidence that stimulus rating [LME - with all trials: t = 1.565, *P* = 0.128, β = 0.045, 95% C.I. = −0.011 – 0.102] was a significant predictor of the participant’s subjective outcome estimation. There was also no statistically significant interaction effect of ‘Rating’ × ‘Incentive Category’ [LME: t = −0.311, *P* = 0.756, β = −0.004, 95% C.I. = −0.028 – 0.021]. In fact, when investigating each category separately, the associations of both curiosity rating for magic tricks [LME: t = 1.618, *P* = 0.118, β = 0.033, 95% C.I. = −0.007 – 0.072] and desirability rating for food items [LME: t = 1.406, *P* = 0.17, β = 0.045, 95% C.I. = −0.018 – 0.108] with the subjective estimation of the outcome were not statistically significant at the set threshold of statistical significance. Comparing the fits of two linear mixed-effects models, one with a ‘stimulus rating’ term and one without, the one without the rating term had a better fit [with all trials, Bayes Factor (BF10) = 0.00559; for magic condition only, BF10 = 0.00572; for food condition only, BF10 = 0.00661; a BF10 value^40^ that is much smaller than 1 indicates very strong support for the null hypothesis compared with the alternative hypothesis including the extra ‘rating’ term, in all cases here]. This means one’s subjective estimation of the chance to win/lose is not likely modulated by the curiosity for information or how desirable the food is.

### Neuroimaging experiments

To examine the neural mechanisms underlying motivational biases on decision-making, we conducted a pair of neuroimaging experiments using fMRI, in which separate groups of participants performed a similar experimental task as in the initial behavioural investigation. The two versions of the fMRI experiment were constructed to be very comparable but differed in the materials used to induce curiosity; an attempt to optimise the generalisability of the findings. The first version (N=31), used magic tricks as curiosity-evoking stimuli (ver1: ‘magic version’) whereas the second version (N=30), used trivia questions (ver2: ‘trivia version’). While magic video clips trigger strong curiosity ^41^, a potential weakness is that they may exert extra demands on visual/perceptual attention and processing for participants, which could be a confound. Trivia questions, on the other hand, have often been used in past experiments of curiosity ^5,6,42,43^, and the visual input is relatively minimal. By running this pair of experiments using different stimuli, we are in a better position to ensure our investigation reflects the impact of curiosity other than some stimulus-specific effect, which has recently been noted as a critical issue in fMRI research ^44^. In addition to the curiosity condition, both versions of the experiment included food trials. We analysed the two versions altogether in order to statistically evaluate the commonalities and differences in neural activation patterns across different types of experimental materials.

Behaviourally, participants from the fMRI experiments (N = 61) accepted the gamble more often when the presented shock probability decreased [GLME: *Z* = −30.357, *P* < 0.001, Exp(β) = 0.337, β = −1.087, 95% C.I.◻=◻−1.157 – −1.017]. Also, on top of the shock probability, participants decided to take the gamble more often when they gave a higher rating to a stimulus in the trial (i.e. higher curiosity/food desirability) [GLME: *Z* = 28.572, *P* < 0.001, Exp (β) = 2.121, β = 0.752, 95% C.I. = 0.700 – 0.804]. Unexpectedly, the fMRI experiments showed a statistically significant interaction between ‘Rating’ and ‘Incentive Category’ [GLME: *Z* = −8.937, *P* < 0.001, Exp (β) = 0.816, β = −0.203, 95% C.I. = −0.248 – −0.159], indicating stronger association of food desirability than curiosity with the acceptance rate in the gambles. Yet it is worth noting that when each category was investigated separately, the link between stimulus rating and acceptance rate in each category was still statistically significant (See Supplementary Table 1). There was also a statistically significant ‘Rating’ × ‘Probability’ interaction effect [GLME: *Z* = −2.786, *P* = 0.0053, Exp (β) = 0.942, β = −0.061, 95% C.I. = −0.103 – −0.018]. Separate analyses for curiosity and food conditions indicated that this interaction effect appeared to be driven mainly by the food condition, which also exhibited an interaction effect between ‘Rating’ and ‘Probability’. Specifically, the impact of food desirability tended to be stronger at lower rates of shock probability [GLME: *Z* = −3.39, *P* <0.001, Exp (β) = 0.905, β = −0.100, 95% C.I. = −0.158 – −0.042]. In contrast, we found no evidence for a statistically significant interaction in the curiosity condition (*P* = 0.397). Since these interaction effects are only specific to these fMRI participants and not consistent with the previous experiments, we will not pursue them further in the current investigation (also see Discussion for a potential interpretation). No other interaction effects reached the set threshold of statistical significance (*Ps* > 0.05). Moreover, an ‘Experiment Version’ effect was also taken into account in this GLME with data from the fMRI participants and we found no evidence that the experiment version has a significant effect on decisions (*P* = 0.662).

Our first fMRI analyses identified brain areas that are recruited at different stages of motivated decision-making and focused on contrasting the trials in which participants accepted the gamble with those that they chose not to (Fig 3a). Based on past studies of extrinsic reward anticipation ^32,45^ and curiosity-guided behavioural change ^5,6^, we hypothesised that activity in the caudate nucleus, NAcc, and VTA/SN would be modulated during the course of curiosity/incentive-driven decision-making. As a number of separate region-of-interest (ROI) masks (one for each of the 3 regions) were tested, we corrected for multiple comparisons across them using a Bonferroni-adjusted alpha level of 0.0167 (familywise error corrected, FWE) per ROI analysis. We focused on the brain activation during the elicitation phase (i.e. when stimuli were presented) and decision phase (i.e. when participants made the decision) in a trial.

**Fig. 3:**
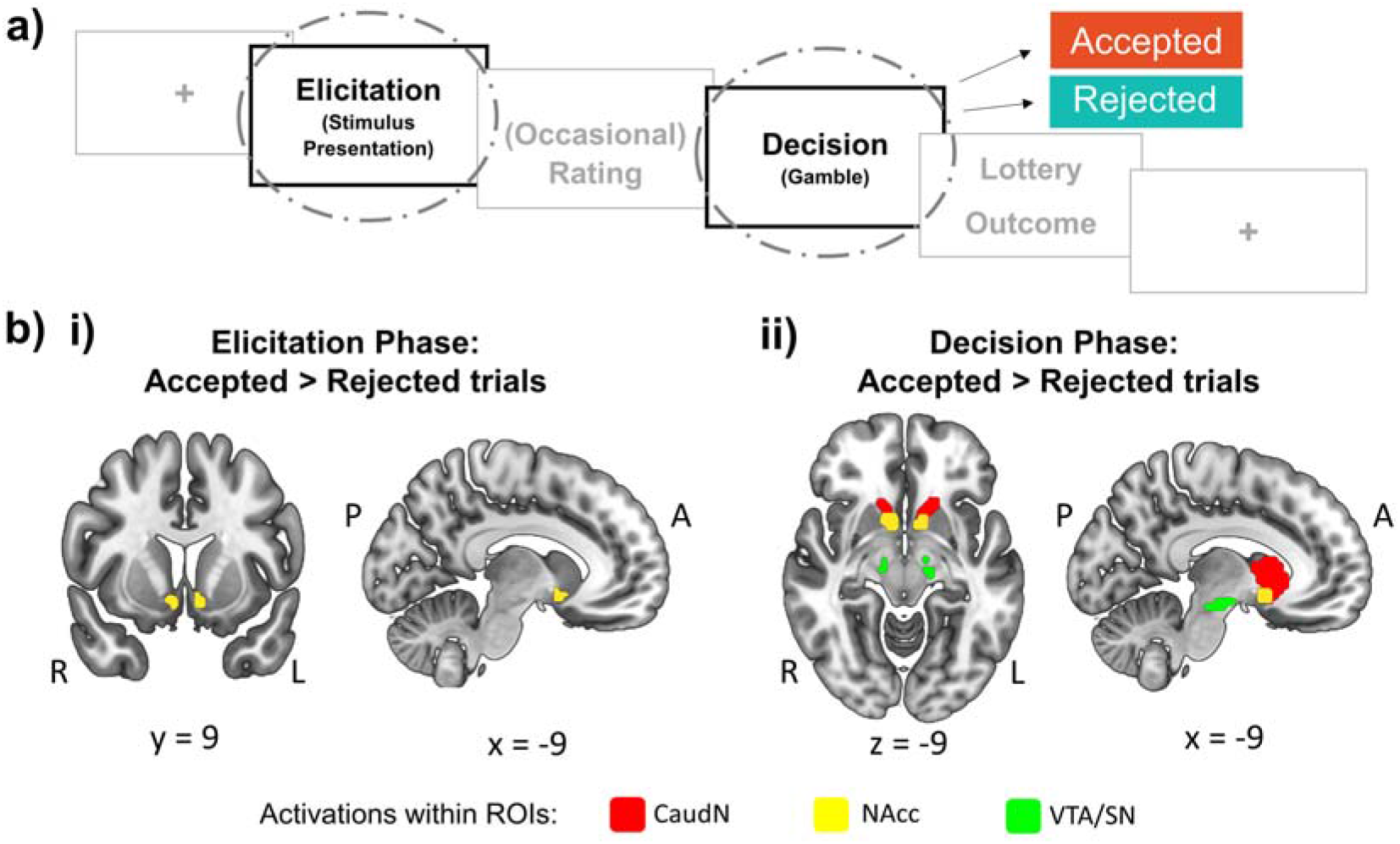
Neural activity in the reward network is modulated by motivation-driven decision-making. a) Brain activity associated with decision-making was analysed according to whether, in the trials, participants accepted or rejected the gamble to satisfy curiosity/obtain the food item at the risk of electric shock. b) Activity was stronger within (i) NAcc at elicitation phase and within (ii) all the dopaminergic circuit ROIs at decision phase for accepted over rejected trials. For visual illustration here, a voxel-wise threshold of P<0.001 (uncorrected) is applied, and all these clusters survived the ROI analysis with an adjusted FWE-corrected statistical significance of P<0.0167 (at cluster level, correcting for multiple comparisons due to the number of ROIs tested). A, anterior; L, left; P, posterior; R, right.

#### Enhanced activity in nucleus accumbens during elicitation of curiosity and food desirability

Comparing all accepted trials with the rejected ones (‘Decision’ contrast) at the elicitation phase revealed enhanced activations specifically in bilateral NAcc (*P_FWE_* < 0.0167, for ROI analysis) among all ROIs (Fig 3b; Supplementary Table 2). There was no statistically significant variation in activation due to the interaction of ‘Decision’ (accept vs. reject) × ‘Experimental Version’ (magic version vs. trivia version), indicating that the type of curiosity-triggering stimuli used tended to have comparable influences on neural responses. On the other hand, a ‘Decision’ × ‘Incentive Category’ (curiosity vs. food) interaction effect was shown, with greater neuronal responses for the ‘accept > reject’ contrast for food compared with the same contrast for curiosity, in a very small cluster of voxels (k=5) in the left NAcc. (Supplementary Table 3).

It is also interesting to observe that while rejected gambles (when compared with accepted ones; i.e., reject > accept ‘Decision’ contrast) did not yield any enhanced responses in the ROIs, an exploratory whole-brain analysis revealed greater activations in the insula and dorsomedial prefrontal cortex encompassing anterior cingulate cortex (*P_FWE_* < 0.05, for whole-brain analysis) (Extended Data Fig 1; Supplementary Table 4), an area previously implicated in value computation^46,47^ and decision conflict^48,49^.

#### Decision to gamble is associated with activity in the brain’s reward network (neural activations at the decision phase)

At the decision phase, a main ‘Decision’ contrast (i.e. accept vs. reject) on our anatomical ROIs revealed that bilateral caudate nucleus, NAcc, and VTA/SN all showed greater activities for the accepted trials in comparison with the rejected trials (*P_FWE_* < 0.0167 for each ROI analysis; see Fig 3b). An exploratory whole-brain analysis of the same contrast also indicated that these striatal and midbrain structures had extensive activations along with other medial brain areas including the cingulate gyrus and medial frontal cortex, as well as the right anterior insula, inferior frontal gyrus and lateral prefrontal cortex (although the peaks were still located in the caudate nucleus) (Extended Data Fig 2; Supplementary Table 5). On the other hand, no statistically significant activation was observed for the reversed contrast (i.e. reject > accept ‘Decision’) at decision phase (Supplementary Table 6). There were also no significant variations in brain activation within the ROIs for the interaction effect of ‘Decision’ × ‘Experimental Version’ (magic version vs. trivia version) or ‘Decision’ × ‘Incentive Category’ (curiosity vs. food) at the decision phase, at the set FWE-corrected statistical threshold. Thus, there is no evidence that the neural responses in the reward system were subject to differential effects due to the type of the curiosity-inducing stimulus and the category of the incentive (see Supplementary Tables 7 & 8).

Next, we examined whether the association between the motivated decision and the striatal activity would still exist after considering the potential effect of the shock (outcome) probability that was shown with the lottery during the decision phase. To this aim, we performed a parametric modulation analysis with another general linear model (GLM), in which the decision to gamble and the presented probability were included as simultaneous parametric modulators for each regressor (i.e. one regressor for curiosity condition; one for food condition) at the decision phase (see Experimental Procedures for details). Indeed, with the presented probability of winning/losing the lottery taken into consideration, the association was still robustly observed in the parametric modulation analysis (despite that the cluster size of activations in each ROI tested has shrunk). Specifically, the activations in bilateral caudate nucleus, NAcc and VTA/SN were shown to be parametrically modulated by the decision of accepting (versus rejecting) to gamble with the ROI approach (*P_FWE_* < 0.0167 in all ROIs; Extended Data Fig. 3), regardless of the incentive category. Moreover, a follow-up whole-brain analysis again revealed extensive neural responses in striatal and midbrain structures, along with the thalamus and right frontal cortex (Supplementary Table 9).

#### Striatal activities mediated the relationship between curiosity/food desirability and the gamble decision

Our behavioural data indicated that the decision to accept/reject a gamble was predicted by the rating of curiosity and food desirability. fMRI results also showed that the decision to accept/reject a gamble was associated with the changes in neural responses in the striatum. Given the literature that the ventral and the dorsal parts of the striatum have been related to different stages of decision-making (the ventral striatum for value computation and the dorsal striatum for action selection ^50–52^; also elucidated in Discussion in more detail), we evaluated whether the link between rating and the decision was mediated by the activities of the ventral striatum (NAcc) at the elicitation phase and dorsal striatum (caudate nucleus) during decision-making on a trial-by-trial basis. The mediation model was tested using a multilevel structural equation model (with trials being nested within participants), and took into account the incentive category and presented shock probability in each trial as well as the experiment version. This multilevel mediation analysis confirmed that stimulus rating predicted choice through the phase-specific activities of nucleus accumbens (at elicitation; Rating ➔ NAcc: *Z* = 3.38, *P* = 0.001, β = 0.08, 95% C.I. = 0.03 – 0.12) and caudate nucleus (at decision-making) (NAcc at elicitation ➔ Caudate at decision: *Z* = 4, *P* < 0.001, β = 0.03, 95% C.I. = 0.02 – 0.05; Caudate at decision ➔ accept/reject response: *Z* = 3.5, *P* < 0.001, β = 0.03, 95% C.I. = 0.01 – 0.04) (see also Fig 4; Supplementary Table 10). The three-path indirect effect (Rating ➔ NAcc at elicitation ➔ Caudate at decision ➔ accept/reject decision response) was statistically significant (*P* = 0.044). These results support the idea that curiosity and food desirability induced by the stimuli prompted participants to make risky decisions through modulating phase-specific striatal activities.

**Fig. 4:**
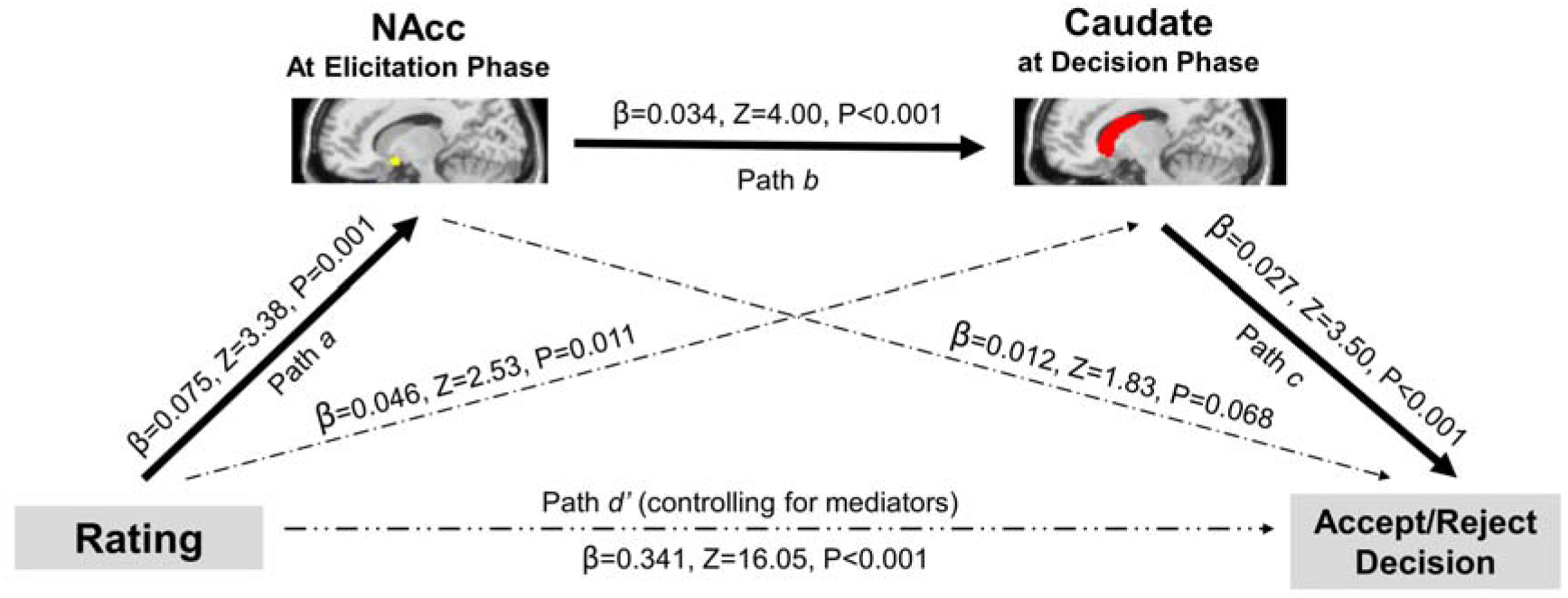
Mediation path diagram. Activations of NAcc and caudate nucleus at different phases of decision-making partially mediated the relationship between the level of curiosity/food desirability (represented by rating) and the decision to gamble. The mediation effect (paths *a* × *b* × *c*) was significant (Z=2.015, P=0.044) and so was the direct path *d*’ (controlling for mediators). The entire analysis also accounted for the incentive category of the stimulus (curiosity-inducing or food), the stimulus version of the experiment (v1 or v2), the presented outcome probability, as well as participants’ characteristics (including gender and subjective shock expectation). Trial-by-trial average striatal activations were extracted within the anatomical masks of bilateral NAcc at the elicitation phase and of bilateral caudate nuclei at the decision phase (see Methods: fMRI analysis). Path lines are labelled with path coefficient (β), Z-value and P-value.

### Exploratory functional connectivity analysis

To further explore the modulation of functional interaction between brain regions during decision-making, we conducted a functional connectivity analysis based on a beta-series correlation approach using the caudate nucleus as the seed. This approach compared correlations of trial-by-trial beta series variability (i.e. variability of series of parameter estimates) between the seed and the rest of the brain in different conditions. We corrected for multiple comparisons across the results using separate left and right caudate as the seed with a Bonferroni-adjusted alpha level of 0.025 per analysis. Using the left caudate nucleus as a seed, this exploratory analysis found weaker connectivity with the left sensorimotor area near precentral gyrus and central sulcus during the decision phase, suggesting a decoupling effect (i.e. weaker functional correlation), when accepting the gamble in comparison with rejecting it. Taking the right caudate nucleus as a seed, we again found weaker connectivity with the left sensorimotor area (a cluster near central sulcus and postcentral gyrus) in accepted (relative to rejected) trials (*P_FWE_* < 0.025 in both analyses; Fig. 5, Supplementary Table 11). The sensorimotor cortex has been associated with virtual feeling and anticipation of pain in action observation ^53–55^. Although speculative, our results might suggest that, when a participant accepted the risk of receiving electric shock, there could be a reduced association between the motivation of satisfying their desire and expected fear of physical shock in the brain. There was no significant variation in brain connectivity shown with the reversed contrast (reject < accept ‘Decision’) and also no significant interaction effects (‘Decision’ × ‘Experiment Version’ or ‘Decision’ × ‘Incentive Category’) at the set FWE-corrected statistical threshold.

**Fig. 5:**
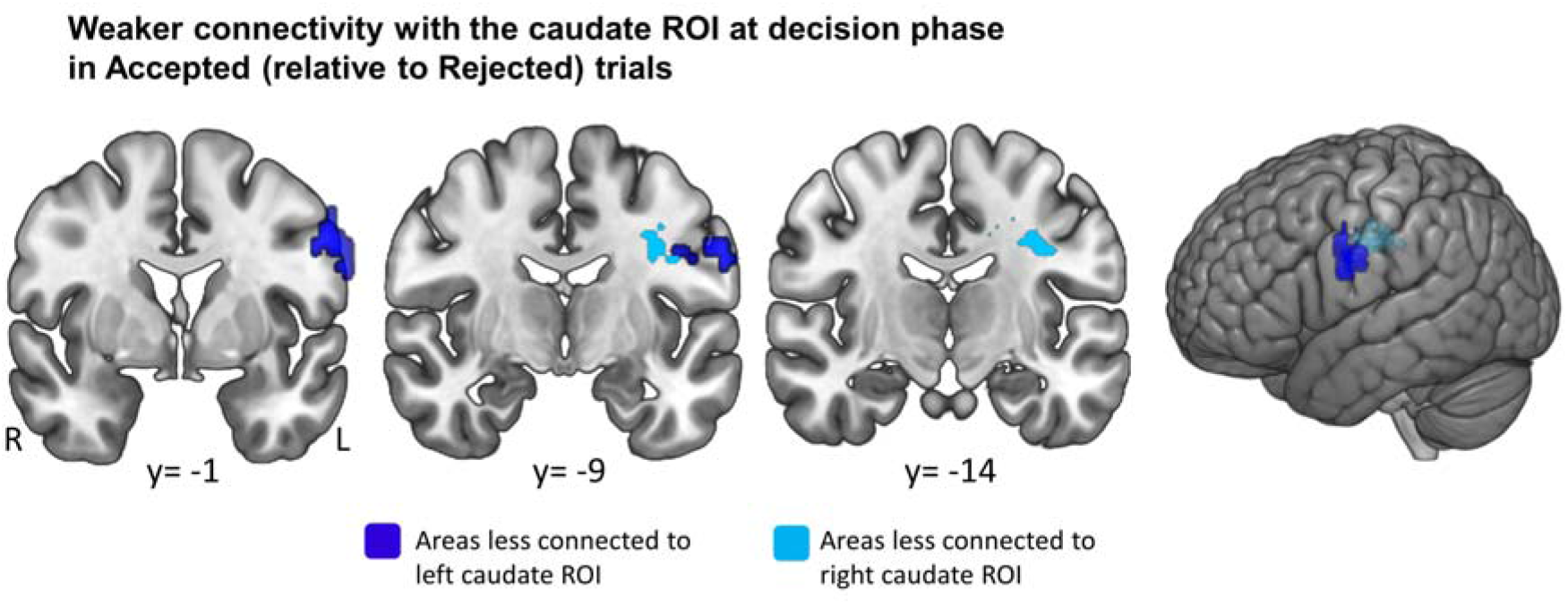
Functional connectivity of caudate nucleus at decision phase. Whole-brain analysis of functional connectivity was based on the beta series correlation approach using anatomical masks of left and right caudate nuclei as the ROIs. A significant main effect of decision was revealed in the left sensorimotor areas (SMA) near central gyrus, exhibiting less connectivity to the left and right caudate ROIs respectively when accepted trials were compared with rejected trials. For visual illustration here, a voxel-wise threshold of p<0.001 (uncorrected) is applied, and all clusters shown survived an adjusted FWE-corrected statistical significance of p<0.025 (correcting for multiple comparisons due to the multiple ROIs applied). L, left; R, right.

## Discussion

Using a curiosity-based decision-making paradigm, the current study aimed to examine whether and how the motivational influence of curiosity overcomes the potential fear of significant risks. Consistent with our speculations, the results showed that i) both curiosity for inconsequential knowledge (magic tricks, trivia questions) and hunger for food indeed prompted participants to subject themselves to physical risks (i.e. electric shocks), and ii) this motivated decision-making was supported by the striatal reward areas. Specifically, the activation in the ventral striatum in response to curiosity-evoking stimuli predicted the decision to accept the risky option through the activation in the dorsal striatum during the decision phase. These results are consistent with the notion that incentive salience plays a role in impulsive behaviours caused by curiosity in a similar manner as it does with extrinsic incentives (e.g., food).

This study extends on the existing neuroscientific literature of curiosity^56^, which primarily focuses on its influence on learning and memory enhancement ^5,6,42^, and our findings provide evidence that corresponds with the idea of a potential role for incentive salience in curiosity-driven risky decision. The influence of curiosity for seemingly useless information in the decision-making process has been recognised in the literature ^57^. Several studies have shown how humans and nonhuman primates have a strong preference for advance information-seeking ^17,18,22,58,59^. That is, if provided with the possibility of receiving advance information about an upcoming reward (e.g. the reward value), individuals have a strong bias to opt to reveal the advance information even if the option incurs a cost (e.g. they have to sacrifice parts of the reward) and also revealing the information in advance itself would not alter the outcome of the reward. Moreover, the motivational lure of curiosity has been demonstrated in recent studies on ‘morbid-curiosity’, where people actively seek negative stimuli (e.g., negative pictures, sounds) in order to satisfy their curiosity for experiencing these uncertain but negative stimuli themselves ^35,60,61^ (although this particular phenomenon may be explained by other motives such as boredom avoidance). The mechanisms underlying curiosity-driven risky decision have been a matter of debate. One of the predominant (but controversial) explanations is that curiosity arouses an aversive state, and it is this aversive state that urges people to acquire information to satisfy their curiosity ^10,11^. However, there has been little empirical evidence that directly supports the assumption that curiosity involves a negative aversive state ^12^ (bar a few exceptions ^62^). Although we did not assess the emotional valence of participants’ feelings of curiosity and therefore cannot directly refute this hypothesis, our findings lend greater support to the incentive salience hypothesis of curiosity, which addresses the strong motivational power of curiosity without making such a restrictive assumption of the aversive nature of curiosity. According to this view, expected acquisition of knowledge, like expected acquisition of food, not only involves the computation of potential expected value of the information, but (irrespective of the expected value of the information) also triggers a strong motivational urge (‘wanting’) that can even overcome the prospective risk of negative consequences to initiate information-seeking behaviour. In other words, underlying the impulsive information seeking behaviour is an active ‘approach’ form of motivation. Another merit of incentive salience hypothesis is that it can be reasonably incorporated into reinforcement learning models of decision-making ^26,63^, providing a more parsimonious and integrative view of the function of curiosity.

Previous fMRI studies of epistemic curiosity using curiosity-inducing materials like trivia questions have reported that the activity of the striatum^5,6^, as well as other areas of the dopaminergic reward network^5^, is enhanced during high states of curiosity when an outcome (i.e. the answer to a trivia question) is anticipated. In addition to the striatal activity during the anticipatory period, one study also found increased activations in the striatum during the “relief of perceptual curiosity” – that is, when a previously blurred image was uncovered and satisfied participants’ curiosity^56^. Our study goes beyond these findings by demonstrating that the striatal responses during anticipatory periods predict actual, motivated choice behaviour (i.e. accepting vs rejecting gambles that entail prospective physical risk). In our study, curiosity for solutions (to magic tricks or trivia questions) is not yet ‘relieved’ in either elicitation or decision phases, and it is reasonable to assume that participants anticipate more to see the solutions in the gambles they accepted than those they rejected, in which they would not anticipate to see the solutions. Consistent with our results, a recent study showed that active choice to reveal uncertain, negative pictures (potentially out of curiosity) is associated with stronger neuronal responses in the striatal areas compared to passive choice of the same pictures^35^. Altogether, the research on the relationship between dopaminergic activity and curiosity states or curiosity-driven behaviours corresponds with prominent curiosity theories that describe curiosity as a motivational state that stimulates active exploration and information seeking^10,14,64^.

Another important finding from our experiments is that curiosity-based decision-making was supported by the different patterns of striatal activation during the elicitation phase (i.e. the ventral striatum) and the decision phase (i.e. broader parts of the striatum peaked at the dorsal striatum); our mediation analysis suggested that the ventral striatum and the dorsal striatum sequentially mediate the relationship between participants’ subjective motivation (i.e. curiosity and hunger for foods) and risky decision-making. These findings dovetail with the notion that the ventral striatum computes the value of the stimulus (‘critic’) while the dorsal striatum plays a role in selecting an action/response (‘actor’) ^50^. Several empirical studies provided supportive evidence for the idea but were all limited to extrinsic incentives ^51,52,65^. Our findings suggest that the neural computation of curiosity-based decision-making can possibly be understood within the actor-critic model of reinforcement learning. In our experiment, the value computation process was potentially initiated when the curiosity-evoking materials were presented (in the elicitation phase) and persevered through the decision phase, during which additional information provided (i.e. outcome probability presented with the wheel of fortune that might affect overall anticipatory value) was incorporated in the overall valuation process; corresponding to this, ventral striatum continued to be recruited throughout these processes. In the meantime, response selection process (whether to approach or avoid information-seeking), potentially supported by the dorsal striatal structures, began in the decision phase. Like many other functional connectivity studies, however, our mediation model is correlational (although there is a temporal separation between the elicitation phase and the decision phase). In addition to this, even within the actor-critic framework, there may be further functional subdivisions of the heterogenous structures of caudate and ventral striatum^66^. Further research with more fine-grained experiments are needed to allow for examination of the potentially dissociable causal roles of various parts of the striatum in curiosity-based decision-making.

In our exploratory analyses, there are some interesting observations that have not been thoroughly discussed in the literature. For example, despite being speculatively only, the decoupling of the dorsal striatum and sensorimotor areas might suggest a neural pathway through which the motivational force of curiosity modulates actual behaviour. The whole-brain fMRI analyses suggested several brain areas related to curiosity-based decision-making that are outside of the dopaminergic system, including the anterior cingulate cortex, lateral prefrontal cortex, and anterior insula. These brain areas have also been reported in previous literature on curiosity ^5,6,35,42,56,67^ (as well as the studies of intrinsically-motivated behaviour in general ^68^). For example, modulated activation in the anterior cingulate cortex and lateral prefrontal cortices has been observed in other whole-brain analyses, elicited by high-compared with low-curiosity trivia questions ^5,42^. Enhanced anterior cingulate and anterior insula activity has also been shown by comparing high with low perceptual uncertainty about an upcoming scene image ^56^, as well as when participants actively choose to reveal negatively valenced information compared with having the information passively assigned to them ^35^. The observed association of curiosity with anterior cingulate cortex and anterior insula, regions sensitive to aversive conditions, may imply an aversive state in curiosity in hindsight. But these regions respond to many other things too, such as surprise (prediction errors ^69,70^) and information update ^67^. These activations may be better understood in terms of a recently proposed framework “Prediction-Appraisal-Curiosity-Exploration” (PACE)^16^ by Gruber and colleagues. This framework suggests that, in the context of curiosity processing, the anterior cingulate cortex serves a role in supporting information-based prediction errors (i.e. information gaps). An information gap is triggered when an event challenges one’s expectations about his/her knowledge on a particular topic. According to PACE framework, when an information gap is recognised, it triggers an appraisal process that is potentially supported by the lateral prefrontal cortex, which then determines one’s actions (e.g. exploration) along with the agent’s subjective experience and the underlying neural mechanism (e.g. curiosity-related dopaminergic processes). In fact, some behavioural studies have highlighted the importance of appraisal processes suggesting that curiosity relies on the appraisal of one’s ability and resources to resolve the challenges raised by the recognition of an information gap ^71–73^. It is worth adding that theories of cognitive control also postulate that anterior cingulate cortex-mediated conflict signals stimulate the lateral prefrontal cortex to direct actions to resolve the conflict ^74,75^. In a dilemma like the one in our experiments that involves prospective physical risk, the appraisal process may also evaluate whether the reward (knowledge acquisition) is worth the cost (risk of shock). If the appraisal determines the information is worth the pursuit, it likely stimulates the exploratory behaviour and information seeking by coupling with the signals from the dopaminergic systems. In keeping with this view, curiosity can be considered as a part of broader autonomous self-regulatory functioning ^15^, which involves the coordination of wide range of brain areas beyond the reward network. For future studies, it would be important to explicitly test and confirm how the different proposed components functionally interact with each other in support of curiosity-based decision.

So far, the existing body of neuroscientific research on curiosity, when interpreting the mechanism of curiosity-driven influence, often attempts to relate to the evidence accumulated on extrinsic incentives, but none of these studies actually compare them directly. In our study, an interesting observation is that we found very few differences overall in brain activation between different types of motivation incentives (i.e. food and curiosity), as far as the decision-making process is concerned (i.e. when comparing accepted with rejected gambles). Consistently, despite the use of different types of materials (i.e. magic trick and trivia questions) to evoke curiosity (with an aim to minimise any stimulus-specific effects), we observed more similarities than differences across the two fMRI experiments. These findings provide strong evidence for the notion that, regardless of the types of incentives, the decision-making process can be portrayed with a common reward-learning framework^13,16,76^. Note that we in fact observed a few differences between the conditions. Food stimuli seemed to have stronger behavioural impacts than curiosity-evoking stimuli in the fMRI experiments (but not in the two behavioural experiments), and this was also reflected in the enhanced activation in a small cluster of voxels in the ventral striatum (NAcc) during the elicitation phase in the food condition than in the curiosity condition. Also, when participants accepted a gamble (vs. rejected it), the magic trick version of the experiment seemed to induce stronger activation in a few voxels in the VTA and SN during the decision phase than the trivia question version; this result suggests a potentially stronger curiosity effect of magic tricks in comparison to trivia questions. However, such differences are all concerned with the magnitude of the motivational effect which is likely to reflect diverse factors irrelevant to how the decision has been made. For example, participants in the fMRI experiments had a prolonged session before the main task actually took place in comparison to those in the two behavioural experiments, due to extra time taken for preparation and mandatory paperwork to undergo an fMRI scan. It is possible that this delay bolstered the hunger of participants (who already had a fasting period before coming to the study), causing stronger effects of hunger both at the behavioural and neural levels. Importantly, however, the incentive salience hypothesis can easily explain the difference in the magnitude of the effects (i.e. food induced stronger incentive salience due to increased hunger) without supposing qualitatively different mechanisms.

Our current findings provide some new insights into the neurocognitive mechanism of how curiosity motivates risky behaviour; however, they should be interpreted with caution. For example, striatal activation has also been implicated in functions other than simple reward processing and incentive salience ^77,78^, thus it is still possible that the observed activation may reflect other decision-making mechanisms than incentive salience. Future studies are needed to examine the neurobiological mechanisms more comprehensively, for example through the use of different methodologies and experimental paradigms such as computational modelling and pharmacological intervention.

## Methods

### Participants

The study was approved by the research ethics committees of the University of Reading, UK (ethics approval number: UREC16/03 & UREC 16/36). Participants were recruited via mailing lists and a research participation pool (managed through SONA Systems) at the University. Participants provided informed consent, completed and passed a health and safety screening on the eligibility of receiving electric stimulation to confirm that i) they did not have a cardiac pacemaker (or any other devices that can be affected by electric stimulation); ii) they were free from neurophysiological symptoms or conditions including peripheral vascular disease, vasculitis cryoglobulinemia, lupus, tingling or numbness in hands and/or feet. To maximise participants’ desire for food during the experiment, they were required not to eat or drink anything (apart from water) within 2 hours prior to the testing session.

Depending on personal preference, participants were compensated either with course credits at a fixed rate of 1 unit per hour or cash payments for their participation. For behavioural experiments, a fixed rate of £7 per hour was given, while for fMRI experiments the compensation for participation was fixed at £10 per hour. In addition, participants were informed that they may receive extra rewards (i.e. food and solutions to magic tricks or trivia questions) according to their task performance in the experiment. Each participant took part in only one version of the experiment.

#### Initial behavioural experiment

Initially, seventeen healthy individuals were recruited; an additional fifteen participants were included later in order to achieve more reliable estimates in data modelling. A final sample of 32 participants (8 males) were 21.90 years old (sd= ±2.66) on average. When analysing these data, we recognise the need to control for the potential inflation of Type-1 error rate resulting from the additional interim analysis performed (see Methods: Behavioural analyses below).

#### Follow-up behavioural experiment (on subjective outcome estimation)

Twenty-nine healthy individuals (7 males) with a mean age of 20.17 years (sd=±3.96) participated in this experiment.

#### fMRI experiments

The first version (magic version) of the experiment included thirty-two individuals. One of them accepted every single gamble (100% acceptance on all trials) in the decision-making task and was thus excluded prior to data analysis. In the second version (trivia version), thirty-two participants were also recruited. One of them accepted every single gamble (100% acceptance on all trials) and another participant had pronounced head movements during MRI scan (> 3mm displacement in a motion direction, see Supplementary Methods: fMRI preprocessing for further details). Both were excluded prior to data analysis. The final sample from both versions comprised a total of 61 participants (11 males) with a mean age of 19.92 years (sd= ± 2.11) and were all right-handed.

For further explanation of the decisions regarding our sample size, see Statement on Statistics and Reproducibility below.

## Materials

### Food images

All pictures were colour photographs selected from different sources on the Internet. They had a resolution of at least 512 × 384 pixels and were edited so that the single food item was presented in the centre against a white background using GNU Image Manipulation Program (GIMP) 2, a free open-source graphics editor. A selection of food was chosen based on the following criteria: first, the items would be familiar to participants to avoid hesitation due to uncertainty during decision-making; second, there was a wide variety of items including fruits/vegetables, sweets, snacks and savoury bites (e.g. grapes, salad, chocolate, nuts, sausage roll, etc.), which might accommodate different individual preferences and tastes for food and elicit different levels of desirability in participants.

### Magic trick videos

Magic tricks, performed by three professional magicians including a champion of an international magic competition, were recorded in a TV studio with a professional cameraman using high resolution video cameras. All videos were then edited using Adobe® Premiere Pro CC® (2015) software to a similar monotonic (dark) background, size (720 × 404 pixels) and viewing focus. The videos were muted (and subtitles were added in a few videos, when needed). The face of the magician was hidden to avoid potential distraction due to appearance and any facial expressions (see Supplementary videos 1-2; refer also to the preprint^79^ from Ozono et al. for more information about this set of magic trick videos). Out of the pool of 166 video clips, we selected magic tricks to be used in the current study by ensuring that they (1) included a range of different features (such as the use of cards, sleight of hands, optical illusions) and (2) likely elicited curiosity to different extents (based on curiosity ratings obtained in a different pilot study). Our initial behavioural experiment included 45 food items and 45 magic trick videos. These videos ranged between 8 – 46 s in length (mean=22.22 s; median=20 s). Due to time constraints (an increase in duration of each trial with an extra event), the follow-up ‘subjective outcome probability estimation’ experiment used only a subset of 36 food items and 36 magic tricks from the initial experiment (length of the videos: range= 8-46 s; mean=20.61 s; median=18.5 s). The magic version of the fMRI experiment used the same subset of stimuli as the follow-up experiment.

### Trivia questions

Sixty trivia questions were selected from a publicly-available 244-item database ^80^ (http://koumurayama.com/resources.php). The selected questions were not obviously culture or age specific. Also, for all questions, the answers were likely to be unknown to the majority of participants. The selection of items corresponded to different trivia categories that might elicit curiosity among individuals to different extents, including art/music, history/geography, movies/TV, nature/animals, science, space, sports, food, as well as other miscellaneous facts. To ensure within-person variability in curiosity evoked across the experiment, half of the chosen questions were picked among those with high mean curiosity scores in the database and the other half among those with low mean scores (rated by a sample of 1498 respondents from a separate online study; for more information, refer to ^80^). The trivia version of the fMRI experiment used 60 trivia questions, as well as 60 food items. On average, the chosen questions contained 10 (ranging between 6-16) words.

#### Experimental Procedure

##### Curiosity-driven Decision-Making Task

The main task of the study followed similar procedures (although there were slight modifications for the follow-up behavioural and fMRI experiments, which are detailed in Supplementary Methods). In brief, each trial started with a central fixation cross and then a brief letter cue (‘M’ signified ‘magic trick’, or ‘T’ for ‘trivia question’ in the ‘trivia version’ fMRI experiment; ‘F’ signified ‘food’) to prepare participants for the kind of stimulus they were about to see. This was followed by the stimulus, which was either a video of a magic trick (or a trivia question in the ‘trivia version’ fMRI experiment) in the curiosity condition or an image of a food item. Participants then gave a rating to indicate their level of curiosity about the magic trick (i.e. how curious they were to see the solution to the trick) or level of desirability of the food (i.e. how much they would like to eat the food), on a 7-point scale (1=not at all, 7=very much). In curiosity trials, they also had to report how confident they were that they knew the solution to a magic trick, using also a 7-point scale (1=not at all, 7=very much). However, because rated confidence was not associated with a participant’s decision and the inclusion of this measure did not change any main results, to make straightforward comparisons between curiosity and food trials, it was not included in the reported analysis here. After rating the stimulus, participants were presented with a wheel of fortune (WoF) representing a lottery which visualised the probability of them winning (the chance of getting the reward) versus losing (the chance of receiving electric shock) in that trial, and were asked to decide whether to gamble or not. Participants were instructed that if they accepted the gamble and won, they would receive a token that might allow them to see the secret behind the magic trick/get the food item after the experiment. If they gambled and lost, they would get a token that might increase the duration of shock they were to experience at the end of the experiment. Participants could also opt to reject the gamble. At the end of each trial, the outcome of the lottery was presented. Participants were informed that the final amounts of rewards and shock to be delivered would be probabilistic as determined by a mathematical algorithm, but with more tokens collected, the likelihood of getting more solutions, food, and shock would be augmented. Participants had an understanding that, as a general rule, the more ‘win’ tokens they collected, the more likely they would get to see more solutions to the tricks and obtain more food items (i.e. the rewards). Similarly, the more ‘loss’ tokens they got, the more likely they would receive more electric shock.

There were 5 versions of WoF, each displaying a different combination of the probabilities of winning and losing a gamble: i) 16.7% (1/6) win vs. 83.3% (5/6) loss; ii) 33.3% (2/6) win vs. 66.7% (4/6) loss; iii) 50% (3/6) win vs. 50% (3/6) loss; iv) 66.7% (4/6) win vs. 33.3% (2/6) loss; v) 83.3% (5/6) win vs. 16.7% (1/6) loss. In the experiments, participants were never shown the actual percentages but the relative win-to-loss contrast of probabilities was illustrated visually by the relative sizes of the constituent slices on the WoF (Fig. 1). To control for the number of ‘success’ and ‘loss’ experiences, unbeknownst to the participants, there was an equal chance of winning or losing in the lottery across all trials.

The curiosity and food trials were mixed and shown in a random order to the participants. The task was programmed and presented using PsychoPy ^81^.

##### Program on the day of the testing session

Participants were asked not to consume any food and drinks (apart from water) within at least 2 hours before attending their testing session, so as to maximise their desire for food during the experiment. This was confirmed with the participants at the beginning of the study, including asking them to indicate when they last ate and had their last meal. Following standard procedures of informed consent and completing corresponding health and safety screening, participants underwent calibration for electric stimulation to identify a maximum (uncomfortable yet non-painful) threshold of electric shock they can endure (see Supplementary Methods for details). Participants did not receive electric shock during the experiment although stimulating electrodes were continuously attached to them. As informed by a pilot study, expectation of electric shock would have more sustained effects on fear perception than receiving actual electric shock multiple times. After the calibration procedures, participants were then given the instructions on the decision-making task (also presented via PsychoPy). There was a practice task that used a different set of stimuli (3 curiosity, 3 food) prior to the main task.

At the end of the experiment, participants were asked in a questionnaire the extent to which they expected to receive electric shock. In this post-experiment questionnaire, the majority of participants across the follow-up behavioural and fMRI experiments reported prospectively that they expected to receive the electric shock during the experiment - 89% of them gave a rating of 3 or above out of 5 (mean=3.67, mode=4), representing their belief that the shock would have happened (we do not have this information from the initial behavioural study). Wherever relevant, in our analysis this shock expectation rating was included as a covariate variable. After completing the questionnaire, rewards (i.e. solutions to some magic tricks and food items) were delivered. Participants did not actually receive any electric shock. They were told that as determined by a probabilistic equation programmed in the task, they were assigned with zero or negligible shock at the end.

#### fMRI acquisition

For participants in the fMRI experiments, whole-brain functional and anatomical images were acquired in a single one-hour scanning session using a 3.0 Tesla Siemens MAGNETOM scanner with a 32-channel Head Matrix coil at the Centre for Integrative Neuroscience and Neurodynamics (CINN), University of Reading.

##### Magic version fMRI experiment

Functional images were acquired using a T2*-weighted gradient-echo echo planar imaging (EPI) pulse sequence with 37 axial slices (in-plane resolution of 3 × 3 × 3mm, interslice gap: 0.75mm), interleaved from bottom to top (echo time (TE): 30 ms; repetition time (TR): 2000 ms; flip angle: 90°; field of view (FOV): 1344 × 1344 mm^2^; in-plane matrix: 64 × 64). A high-resolution T1-weighted three-dimensional anatomical image was also collected, using an MPRAGE-gradient sequence with 176 × 1mm slices (in-plane resolution of 1 × 1 × 1 mm; TE: 2.52 ms; TR: 2020 ms; Inversion Time (TI):1100 ms; FOV: 250 × 250; flip angle: 9°), enabling optimal localisation of the functional effects.

##### Trivia version fMRI experiment

Functional images were acquired using the same sequence and parameters as in fMRI experiment 1. A high-resolution T1-weighted three-dimensional anatomical image was collected using an MPRAGE-gradient sequence with 192 × 1 mm slices (in-plane resolution of 1 × 1 × 1 mm; TE: 2.29 ms; TR: 2300 ms; TI:900 ms; FOV: 240 × 240; flip angle: 8°).

#### fMRI analysis

Preprocessing and data analyses of the imaging data were performed using the SPM12 software (www.fil.ion.ucl.ac.uk/~spm). The preprocessing procedures included spatial realignment of the EPI volumes, co-registration with the structural image, segmentation, group-wise normalisation using DARTEL, and smoothing (see Supplementary Methods for more details).

##### Region-of-Interest (ROI) Mask

Based on past findings, the striatal reward network is modulated by motivational states and plays an important role in influencing motivation-driven behaviour ^5,6,82,83^. We examined bilateral striatal structures including caudate nucleus and NAcc, as well as VTA/SN, and performed in these anatomical masks the small volume correction ^84^, implemented in SPM. The masks were anatomically defined using a high-resolution probabilistic atlas of human subcortical brain nuclei by Pauli and colleagues ^85^, and only the voxels that were part of the brain areas of interest with a probability of at least 15% were included in the ROIs. Our ROI masks included 565 voxels (3.7 resels) for caudate nucleus, 53 voxels (0.3 resels) for NAcc, and 66 voxels (0.3 resels) for VTA/SN. Results were yielded with a height-defining threshold at voxel-level *P* < 0.001 and a cluster extent: k ≥ 5. Also, to control for the risk of type I error due to multiple comparisons using several anatomical masks, we focused on results that only survived an adjusted familywise error-corrected significance threshold of *P* < 0.0167 at cluster level (i.e. 0.05 over 3 comparisons).

As an exploratory attempt to examine any other brain regions outside of the reward system that are related to motivation-driven decisions, in addition to the ROI approach, we also analysed the fMRI data with a whole-brain approach. Again, results were obtained using a height-defining threshold at voxel-level *P* < 0.001, and we focused only on results that survived a familywise error-corrected significance threshold of *P* < 0.05 at cluster level.

##### GLM - Activation predicting motivated decision-making

One of the aims of the current study was to test whether curiosity about knowledge and desire for food influenced a participant’s decision in a similar manner via the reward system and also whether there were similarities and differences in effects depending on the type of stimuli presented. To this aim, we implemented a general linear model (GLM 1) for each subject that regressed brain activation depending on the dichotomous decision in the lottery – whether an individual accepted or rejected it – in each incentive category. Specifically, four separate regressors were specified for the accepted and rejected trials in curiosity and food categories (i.e. 4 conditions), at the onset of the decision phase of each trial (i.e. when a wheel of fortune appeared and the participant had to make a choice). To account for the brain activation related to the elicitation of curiosity and food desirability, another 4 regressors were specified to model the stimulus presentation onset (for magic trick trials specifically, this was time-locked to the moment of subjective surprise defined by the group; see Supplementary Methods for details) of the accepted and rejected trials in the two categories. Moreover, reaction time of the decision response was added for parametric modulation in the four ‘decision phase’ regressors and head motion parameters were included as regressors of no interest. To test for group-level effects, beta images for the four conditions from each subject in the fMRI experiments were entered as input data for second-level analysis. Using a flexible factorial design in the second-level models (separately for the elicitation phase and the decision phase), we specified the factors of ‘Decision’ (accept vs. reject), ‘Incentive Category’ (food vs. curiosity) and ‘Experimental Version’ (magic vs. trivia), and included each subject’s gender and shock expectation rating (see above) as covariates of no interest. Our main focus was the main effect of ‘Decision’ (i.e. accepted vs. rejected trials) to examine if curiosity and food influenced decision-making in a similar way via the reward system. We also examined the interaction effects to probe any potential variations in neural activity specific to the type of materials used (i.e. stimuli of different incentive categories or different curiosity-triggering materials).

To run a multilevel mediation analysis (explained below), we also created a different linear model (GLM 2) in which each trial was modelled as a separate regressor. These models (implemented separately for the elicitation phase and the decision phase) enabled us to estimate a separate statistical map for each trial (i.e. single-trial activation patterns). We were then able to extract parameter estimates (beta values associated with ‘brain activations’) for each trial in these maps as inputs for the mediation analysis. As was the case in GLM 1, head motion parameters were included as regressors of no interest.

##### Parametric modulation analysis - Accounting for the presented outcome probability in evaluating the relationship between decision and brain activation

To account for the effect of the presented probability of winning/losing on decisions, we implemented a further linear model (GLM 3) and performed parametric modulation analysis. This time, in first-level design, two regressors were specified to model the onsets of the decision phase of trials for the two incentive categories (curiosity, food). To each regressor, the participant’s decision in the lottery was added as a parametric modulator (PM) (i.e. ‘accept’ choice was coded as +1; ‘reject’ choice as −1); the corresponding outcome probability presented in each trial was also centred and added as another PM (i.e. 83.3% shock probability was coded as −2; 66.7% as −1; 50% as 0; 33.3% as 1; and 16.7% as 2). In addition, we included reaction time as another PM. Importantly, these parametric modulators were not orthogonalized, meaning that no priority was given specifically to the first PM (or a particular PM) over other PMs in explaining the variance in neural response (see the method paper by Mumford and colleagues ^86^). The onsets of the elicitation phase (stimulus presentation) for the two incentive categories were specified as separate regressors, along with the head motion parameters as regressors of no interest. Our main focus of interest here was to examine whether the ‘accept/reject’ decision (i.e. the ‘Decision’ PM) still parametrically modulated activations in the pre-defined ROIs even after taking the presented outcome probability of into account.

##### Functional Connectivity Analysis

Functional connectivity was examined with the beta series correlation method ^87^ implemented in BASCO toolbox ^88^. This method allowed us to use trial-by-trial variability to characterise dynamic inter-regional interactions. We specifically tried to explore the functional interaction between brain regions during decision-making and used the left and right caudate nuclei as the ROIs in this connectivity analysis (as the peaked activations at the decision phase were shown within the caudate in the main fMRI analysis; see Results). The anatomical masks used here were defined similarly as in the ROI approach of the main fMRI analysis above. Left and right caudate was analysed separately because brain connectivity might be strongly constrained by laterality ^89,90^. We corrected for multiple comparisons across the results using separate left and right caudate seeds with a Bonferroni-adjusted (FWE-corrected) alpha level of 0.025 per analysis.

At the first level of the analysis, a new GLM was constructed (implemented in BASCO), in which BOLD response time-locked to the onset of the decision phase of each trial was modelled individually by a separate regressor using a canonical haemodynamic response function. This resulted in different parameter estimates for each trial for each participant. The six motion parameters for each run were also included in this GLM. Next, seed-based correlations were computed voxel-wise for each participant and for each of the experimental conditions of interest. This procedure generated an individual’s correlation map between each seed region’s beta series and the beta series of all other voxels in the brain separately for each condition of interest, which was normalised using Fisher’s r-to-z transformation. At the second level, the set of correlation maps from each participant was subjected to random-effects analysis to identify voxels that showed changes in functional connectivity with the seed (based on trial-by-trial variability in parameter estimates) across different conditions at the group level. Specifically, we used a flexible factorial design (implemented in SPM) and specified factors of ‘Decision’ as well as ‘Incentive Category’ and ‘Experiment Version’. We also included the participant’s gender and his/her shock expectation rating as covariates of no interest. Again, our main focus was the main effect of ‘Decision’ (i.e. accepted vs. rejected trials), but we also examined the interaction effects to probe any potential variations in functional connectivity that may be specific to the type of materials used (i.e. stimuli of different incentive categories or different curiosity-triggering materials).

#### Behavioural analysis

##### Linear mixed-effects modelling

To examine the main question of whether curiosity and food desirability influence decisions on a trial-by-trial basis at the behavioural level, we performed a generalised linear mixed-effects modelling (GLME) analysis. In the ‘full’ model, a participant’s decision in the lottery (i.e. accept or reject) was specified as the dichotomous outcome, in a logistic link function. Stimulus rating (group-mean centred), the presented shock probability, and the category of incentive (curiosity vs. food) as well as their interaction terms (including the three-way interaction), were entered as predictors of the decision. To account for the nested structure of the data, we specified random intercepts and slopes of participants.

Mixed-effects model with multiple random slopes is prone to convergence errors given the complexity of the model ^91^. We first modelled all the possible random slopes and then dropped one random slope each time until model convergence (i.e. error-free). The results reported here were based on the models acquired with this strategy. Importantly, we also carried out a sensitivity check to make sure that the results were not biased by omitting specific random slopes ^92^. The sensitivity analysis also check for the random effects of stimulus as this effect could potentially inflate Type-1 error rates ^93^. All the reported results were demonstrated to be very robust to the specification of random effects.

We analysed behavioural data from the initial experiment, the follow-up experiment and the fMRI experiments. In the GLME for the fMRI experiments, we added the experiment version (ver 1: magic vs. ver 2: trivia) as an extra fixed-effect term. Note that this ‘experiment version’ term was not a significant predictor, indicating no evidence for differences in participants’ performance (in terms of acceptance rate) between different versions of the experiment. To facilitate the understanding of the results further, Supplementary Table 1 present GLME results separately for curiosity and food conditions.

As noted earlier, for the initial behavioural experiment we carried out additional data collection in order to attain more reliable parameter estimates. We recognise this procedure would result in repeated analyses (as data accumulation continued), thereby increasing the potential risk of overestimation of any effects and type 1 error ^94^. To control for this risk, we followed a conservative Šidák approach^94,95^ to adjust critical P-value for multiple interim analyses conducted (similar to multiple comparisons). Based on the Šidák adjustment [P’ = 1− (1− P)^1/k^], with an ordinary critical P-value being 0.05 and the number of interim analyses k=2 in our case, the adjusted critical value P’ would be 0.025. Any analyses reported here that used data from the initial behavioural experiment applied this adjusted critical value as a cut-off in significance testing.

The follow-up ‘subjective outcome estimation’ experiment had a specific aim to investigate whether curiosity and food desirability distort a participant’s subjective estimation of the probability of winning/losing the lottery (a continuous dependent variable). The main analysis in this experiment was based on a linear mixed-effects model (LME) that included the stimulus rating (curiosity/food desirability) and the presented probability of winning/losing as predictors of subjective outcome probability estimation. Again, we specified random intercepts and slopes of participants. We compared two models, one including the stimulus rating and one without, and computed Bayes factors based on the Bayesian Information Criterion (BIC) from each model (readily extracted using ‘BIC’ function in R) to evaluate the effect of the rating. A measure of BIC quantifies a formal model’s goodness of fit to data, taking into account the number of free parameters in the model ^96,97^. A lower BIC value indicates a better fit, and vice versa. As described by Wagenmakers^97^, Bayes factor (BF) can be estimated using this transformation (exponentiation of the average of the difference in BIC values for two competing models, i & j): BF ≃ exp[(BIC_i_ − BIC_j_)/2] (see also Masson’s tutorial ^98^). The resulting estimate of the Bayes factor yields the odds favouring the alternative hypothesis, relative to the null hypothesis.

Testing of GLME and LME models was carried out using the package ‘lme4’ ^99^ and graph plotting of the behavioural data was achieved using ‘ggplot’ and ‘raincloud plot’ ^100^ in R.

##### Mediation analysis

To test whether the link between rated curiosity/food desirability and the decision to gamble is mediated by the striatal activations observed in the main fMRI analysis, we performed a multilevel mediation analysis using multilevel structural equation modelling with Mplus ^101^ (version 7). Specifically, we attempted to test the involvement of the activities of the ventral striatum during the elicitation phase and the dorsal striatum during decision-making in mediating the link.

In this analysis, we specified the indirect paths as starting from Rating through Striatal activations at elicitation phase and decision phase to Decision response (i.e. Rating ➔ NAcc activity at Elicitation ➔ Caudate activity at Decision-Making ➔ Decision response) as well as the corresponding direct paths, and analysed data of curiosity and food conditions altogether using trials as the unit of analysis (Fig 4). Incentive category (curiosity vs. food), the presented shock probability, as well as the experiment version were included as covariates in the model. Moreover, gender of the participant as well as their shock expectation rating in the post-experiment questionnaire were accounted for in the model as subject-level covariates. Decision here is a binary outcome variable (accept vs. reject).

To account for the multilevel nature of the data, we used person-mean centring ^102^ before parameter estimation and computed cluster robust-standard error to make statistical inference ^103^. After estimating the model, the standard error of the mediation effect was computed using the delta method and statistical significance of the mediation effect was tested.

#### Statement on statistics and reproducibility

The study employed a within-subject design and participants were not assigned to different experimental treatments. Data collection and analysis were not performed blind to the conditions of the experiments. All statistical tests on behavioural data were two-tailed and used an alpha level of 0.05 (except analysis with data from the initial behavioural experiment and unless otherwise stated). In the analysis with continuous dependent variable, residual distribution was assumed to be normal but this was not formally tested. By making reference to the estimated effect size of a separate pilot study (N = 34), the original sample size of the initial behavioural experiment (N = 17) was sufficient to detect the estimated effect at a statistical power of 95%. Additional fifteen participants were added later into this initial experiment in order to achieve more reliable estimates in data modelling and also for better comparison with other experiments (upon reviewers’ suggestion). To control for the potential risk of inflated type 1 error due to this subsequent addition of participants, we followed the Šidák approach^94,95^ to adjust critical P-value for multiple interim analyses conducted (see Methods above). For the follow-up behavioural experiment, the sample size (N = 29) was sufficient to detect a small to medium effect (of curiosity rating on subjective probability estimation) with a statistical power of 80% ^104^. The sample sizes of the fMRI experiments were determined to be comparable to or larger than other neuroimaging studies of motivational biases ^105^ and curiosity-based effects ^5,6^. Also, such a sample size should be sufficient to reliably detect medium effect sizes in task-based fMRI analysis ^106^.

## Data availability

The behavioural data that support the findings of the current study would be available at https://osf.io/mafe3/. The unthresholded statistical maps of the fMRI results can be accessed at https://neurovault.org/collections/AWZZIZCZ/.

## Code availability

The analyses in this study were performed in standard software and based on published routine, as specified in detail in the Methods and the Supplementary Information. Custom codes can be accessed through https://osf.io/mafe3/ and are available from the corresponding authors on request.

## Acknowledgments

The study was supported by the Marie Curie Career Integration Grant (CIG630680 to K.M.), JSPS KAKENHI (15H05401; 16H06406, 18H01102, and 18K18696 to K.M.), F. J. McGuigan Early Career Investigator Prize (to K.M.), Jacobs Foundation Advanced Fellowship (to K.M.) and the Leverhulme Trust (RPG-2016-146 and RL-2016-030 to K.M.). The funders had no role in study design, data collection and analysis, decision to publish or preparation of the manuscript. We are grateful of the magicians (including Shota Irieda, Ohkubo Kohei and Malka) for producing magic tricks for our research. We thank C. Ogulmus, E. Daveau and the rest of the MeMo Lab as well as S. Shen and the CINN for helping with data collection, C. Inaltay, A. Firat, A. Haffey, J. Raw, and G. Fastrich for editing and pilot-testing magic video clips, A. Mihalik for advice on further advanced analysis, and C. McNabb for providing useful comments on the drafts of the article.

## Author contributions

K.M. conceived the idea; J.K.L.L. and K.M. jointly designed the study; J.K.L.L., H.O., A.K., and K.K. created experimental materials. J.K.L.L. performed research and analysed the data; J.K.L.L. and K.M. jointly wrote the paper. All authors provided critical comments.

---

**Extended Data Fig 1.**
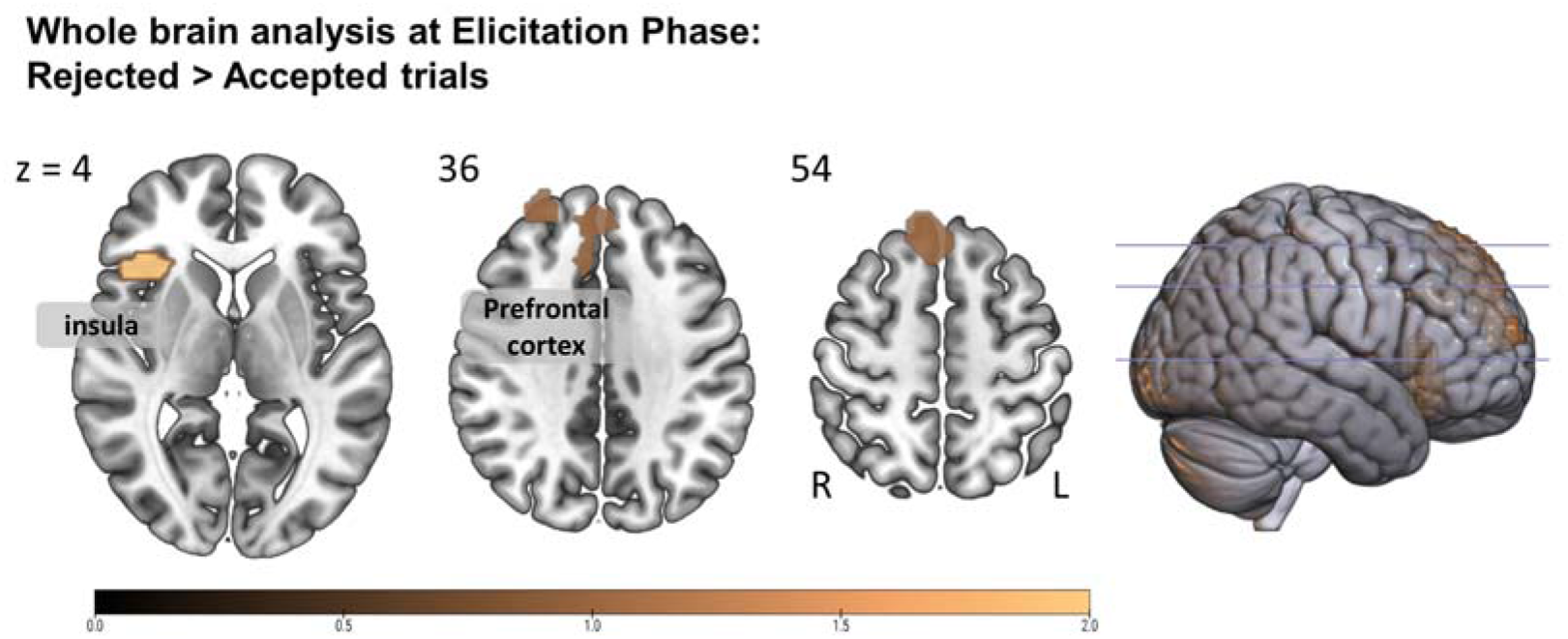
Exploratory whole-brain analyses at elicitation phase showed differential brain activations when comparing ‘rejected’ with ‘accepted’ trials. Activity was stronger within prefrontal cortex and insular gyrus at the elicitation phase of trials in which participants rejected the gamble. For visual illustration here, a voxel-wise threshold of P<0.001 (uncorrected) is applied; all clusters survived a FWE-corrected statistical significance threshold of P<0.05 (at cluster level). L, left; R, right.

**Extended Data Fig 2.**
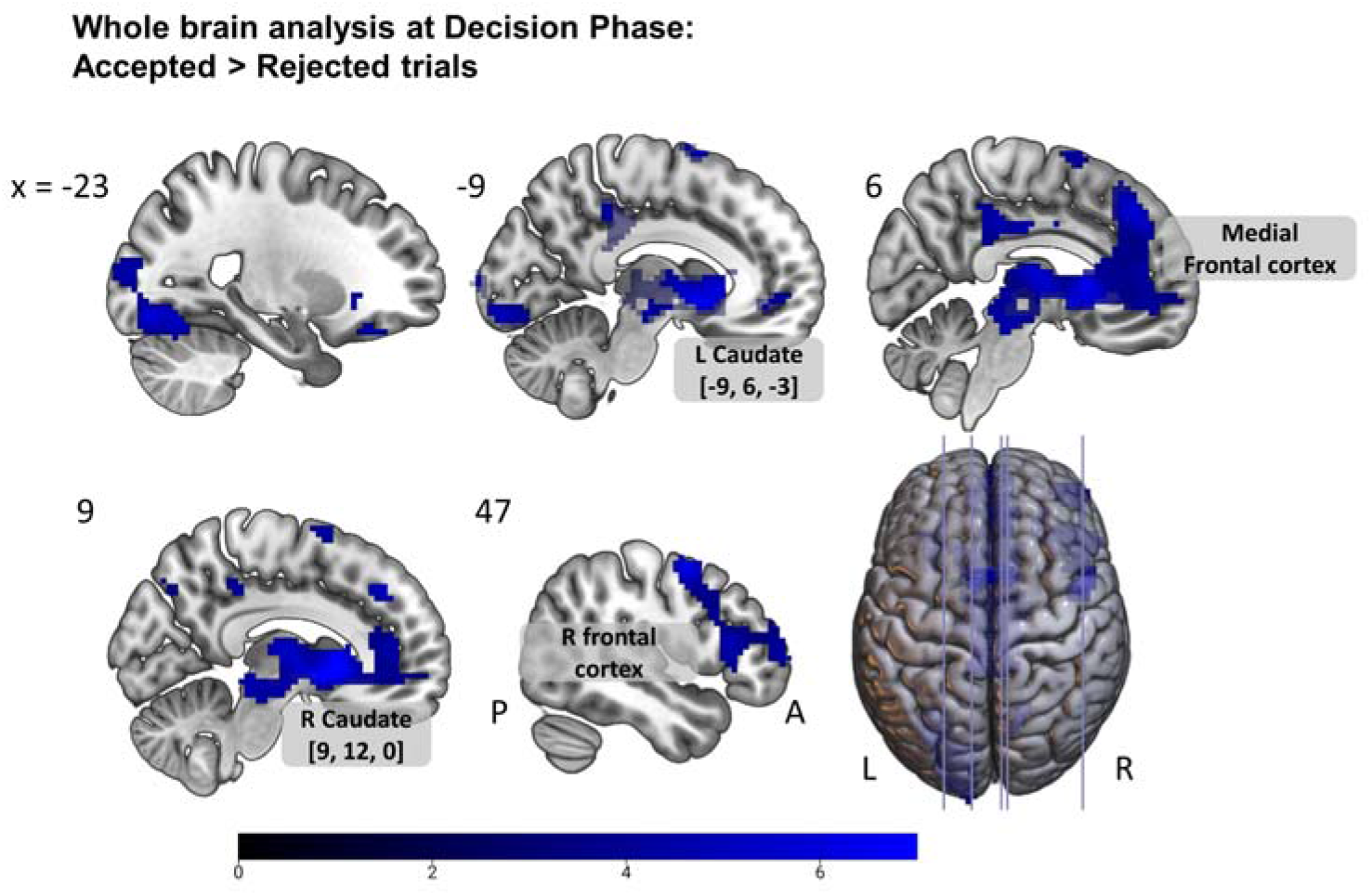
Exploratory whole-brain analyses with a main effect of decision (accepted > rejected trials) at decision phase. Peak activation is shown for the right caudate nucleus (MNI coordinate: 9, 12, 0) and the left caudate nucleus (MNI coordinate: −9, 6, −3) in an extensive medial reward network cluster, extending into the thalamus and the medial frontal cortex, as well as the right frontal cortex and anterior insula. For visual illustration here, a voxel-wise threshold of P<0.001 (uncorrected) is applied; all clusters survived a FWE-corrected statistical significance threshold of P<0.05 (at cluster level). See ROI results in Figure 3. A, anterior; L, left; P, posterior; R, right.

**Extended Data Fig 3.**
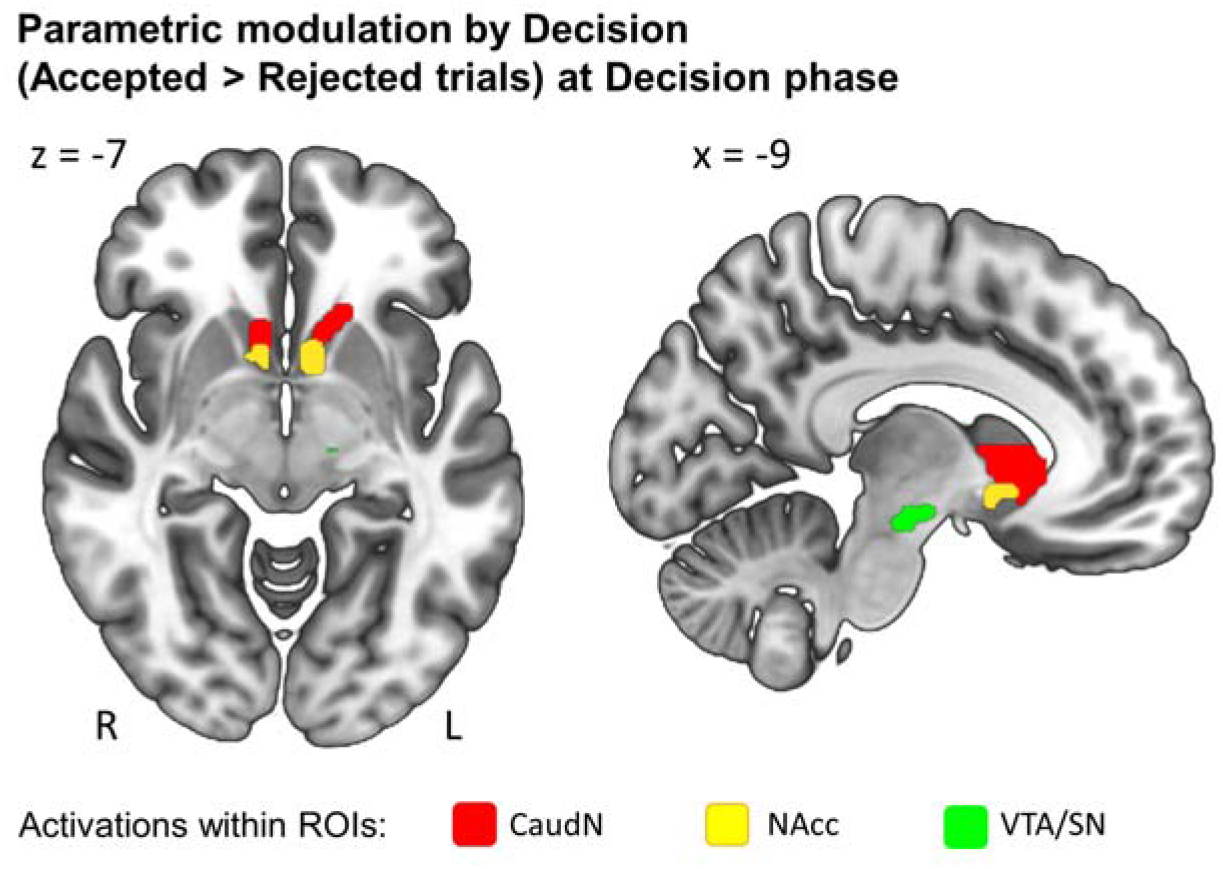
ROI activations for motivation-driven decision-making in a parametric modulation analysis accounting for presented outcome probability. Differential activations for accepted (> rejected) trials were observed within the ROIs of caudate nucleus, NAcc, and VTA/SN at the Decision phase, even when taking into account the shock/outcome probability presented as an additional parametric modulator in the model. For visual illustration, a voxel-wise threshold of P<0.001 (uncorrected) is applied here; clusters survived the ROI analysis with an adjusted FWE-corrected statistical significance threshold of P<0.0167 (at cluster level). A, anterior; L, left; P, posterior; R, right.

## References

1. Gottlieb, J., Oudeyer, P.-Y., Lopes, M. & Baranes, A. Information-seeking, curiosity, and attention: computational and neural mechanisms. Trends Cogn. Sci. 17, 585–593 (2013).

2. Jirout, J. & Klahr, D. Children’s scientific curiosity: In search of an operational definition of an elusive concept. Dev. Rev. 32, 125–160 (2012).

3. Kidd, C. & Hayden, B. Y. The psychology and neuroscience of curiosity. Neuron 88, 449–460 (2015).

4. von Stumm, S., Hell, B. & Chamorro-Premuzic, T. The hungry mind: Intellectual curiosity is the third pillar of academic performance. Perspect. Psychol. Sci. 6, 574–588 (2011).

5. Gruber, M. J., Gelman, B. D. & Ranganath, C. States of curiosity modulate hippocampus-dependent learning via the dopaminergic circuit. Neuron 84, 486–496 (2014).

6. Kang, M. J. et al. The wick in the candle of learning: Epistemic curiosity activates reward circuitry and enhances memory. Psychol. Sci. 20, 963–973 (2009).

7. Renninger, K. A. & Hidi, S. The power of interest for motivation and engagement. (Routledge, NY, 2016).

8. Sakaki, M., Yagi, A. & Murayama, K. Curiosity in old age: A possible key to achieving adaptive aging. Neurosci. Biobehav. Rev. 88, 106–116 (2018).

9. Rozek, C. S., Svoboda, R. C., Harackiewicz, J. M., Hulleman, C. S. & Hyde, J. S. Utility-value intervention with parents increases students’ STEM preparation and career pursuit. Proc. Natl. Acad. Sci. 114, 909–914 (2017).

10. Loewenstein, G. The psychology of curiosity: A review and reinterpretation. Psychol. Bull. 116, 75–98 (1994).

11. Berlyne, D. E. Conflict, arousal, and curiosity. (McGraw-Hill Book Company, 1960). doi:10.1037/11164-000

12. Silvia, P. J. Exploring the Psychology of Interest. (Oxford University Press, 2006). doi:10.1093/acprof:oso/9780195158557.001.0001

13. Marvin, C. B. & Shohamy, D. Curiosity and reward: Valence predicts choice and information prediction errors enhance learning. J. Exp. Psychol. Gen. 145, 266–272 (2016).

14. Gottlieb, J. & Oudeyer, P.-Y. Towards a neuroscience of active sampling and curiosity. Nat. Rev. Neurosci. 19, 758–770 (2018).

15. Murayama, K. A reward-learning framework of autonomous knowledge acquisition: An integrated account of curiosity, interest, and intrinsic-extrinsic rewards. Preprint at OSF doi:10.31219/osf.io/zey4k (2019).

16. Gruber, M. J. & Ranganath, C. How Curiosity Enhances Hippocampus-Dependent Memory: The Prediction, Appraisal, Curiosity, and Exploration (PACE) Framework. Trends Cogn. Sci. 23, 1014–1025 (2019).

17. Kobayashi, K., Ravaioli, S., Baranès, A., Woodford, M. & Gottlieb, J. Diverse motives for human curiosity. Nat. Hum. Behav. 3, 587–595 (2019).

18. Blanchard, T. C., Hayden, B. Y. & Bromberg-Martin, E. S. Orbitofrontal cortex uses distinct codes for different choice attributes in decisions motivated by curiosity. Neuron 85, 602–614 (2015).

19. Daddaoua, N., Lopes, M. & Gottlieb, J. Intrinsically motivated oculomotor exploration guided by uncertainty reduction and conditioned reinforcement in non-human primates. Sci. Rep. 6, 20202 (2016).

20. Vasconcelos, M., Monteiro, T. & Kacelnik, A. Irrational choice and the value of information. Sci. Rep. 5, 13874 (2015).

21. Bennett, D., Bode, S., Brydevall, M., Warren, H. & Murawski, C. Intrinsic valuation of information in decision making under uncertainty. PLOS Comput. Biol. 12, e1005020 (2016).

22. Brydevall, M., Bennett, D., Murawski, C. & Bode, S. The neural encoding of information prediction errors during non-instrumental information seeking. Sci. Rep. 8, 6134 (2018).

23. Eliaz, K. & Schotter, A. Paying for confidence: An experimental study of the demand for non-instrumental information. Games Econ. Behav. 70, 304–324 (2010).

24. Berridge, K. C. Motivation concepts in behavioral neuroscience. Physiol. Behav. 81, 179–209 (2004).

25. Anselme, P. & Robinson, M. J. F. Incentive Motivation: The Missing Piece between Learning and Behavior. in The Cambridge Handbook of Motivation and Learning (eds. Renninger, K. A. & Hidi, S.) 163–182 (Cambridge University Press, 2019). doi:10.1017/9781316823279.009

26. Berridge, K. C. From prediction error to incentive salience: mesolimbic computation of reward motivation. Eur. J. Neurosci. 35, 1124–1143 (2012).

27. Robinson, T. E. & Berridge, K. C. The incentive sensitization theory of addiction: some current issues. Philos. Trans. R. Soc. B Biol. Sci. 363, 3137–3146 (2008).

28. Kringelbach, M. L. & Berridge, K. C. Neuroscience of Reward, Motivation, and Drive. in Recent Developments in Neuroscience Research on Human Motivation (eds. Kim, S., Reeve, J. & Bong, M.) 23–35 (Emerald Group Publishing Limited, Bingley, 2016). doi:10.1108/S0749-742320160000019020

29. Berridge, K. C. ‘Liking’ and ‘wanting’ food rewards: Brain substrates and roles in eating disorders. Physiol. Behav. 97, 537–550 (2009).

30. Tang, D. W., Fellows, L. K., Small, D. M. & Dagher, A. Food and drug cues activate similar brain regions: A meta-analysis of functional MRI studies. Physiol. Behav. 106, 317–324 (2012).

31. O’Doherty, J. P., Deichmann, R., Critchley, H. D. & Dolan, R. J. Neural Responses during Anticipation of a Primary Taste Reward. Neuron 33, 815–826 (2002).

32. Knutson, B., Adams, C. M., Fong, G. W. & Hommer, D. Anticipation of Increasing Monetary Reward Selectively Recruits Nucleus Accumbens. J. Neurosci. 21, RC159–RC159 (2001).

33. Lawrence, N. S., Hinton, E. C., Parkinson, J. A. & Lawrence, A. D. Nucleus accumbens response to food cues predicts subsequent snack consumption in women and increased body mass index in those with reduced self-control. Neuroimage 63, 415–422 (2012).

34. Litman, J. Curiosity and the pleasures of learning: Wanting and liking new information. Cogn. Emot. 19, 793–814 (2005).

35. Oosterwijk, S., Snoek, L., Tekoppele, J., Engelbert, L. & Scholte, H. S. Choosing to view morbid information involves reward circuitry. Preprint at bioRxiv doi:10.1101/795120 (2019).

36. Kobayashi, K. & Hsu, M. Common neural code for reward and information value. Proc. Natl. Acad. Sci. 116, 13061–13066 (2019).

37. Gerdeman, G. L., Partridge, J. G., Lupica, C. R. & Lovinger, D. M. It could be habit forming: drugs of abuse and striatal synaptic plasticity. Trends Neurosci. 26, 184–192 (2003).

38. Telzer, E. H. Dopaminergic reward sensitivity can promote adolescent health: A new perspective on the mechanism of ventral striatum activation. Dev. Cogn. Neurosci. 17, 57–67 (2016).

39. Wright, W. F. & Bower, G. H. Mood effects on subjective probability assessment. Organ. Behav. Hum. Decis. Process. 52, 276–291 (1992).

40. van Doorn, J. et al. The JASP Guidelines for Conducting and Reporting a Bayesian Analysis. Preprint at PsyArxiv doi:10.31234/osf.io/yqxfr (2019).

41. Moss, S. A., Irons, M. & Boland, M. The magic of magic: The effect of magic tricks on subsequent engagement with lecture material. Br. J. Educ. Psychol. 87, 32–42 (2017).

42. Ligneul, R., Mermillod, M. & Morisseau, T. From relief to surprise: Dual control of epistemic curiosity in the human brain. Neuroimage 181, 490–500 (2018).

43. Baranes, A., Oudeyer, P.-Y. & Gottlieb, J. Eye movements reveal epistemic curiosity in human observers. Vision Res. 117, 81–90 (2015).

44. Westfall, J., Nichols, T. E. & Yarkoni, T. Fixing the stimulus-as-fixed-effect fallacy in task fMRI. Wellcome Open Res. 1, 23 (2016).

45. Adcock, R. A., Thangavel, A., Whitfield-Gabrieli, S., Knutson, B. & Gabrieli, J. D. E. Reward-Motivated Learning: Mesolimbic Activation Precedes Memory Formation. Neuron 50, 507–517 (2006).

46. Shenhav, A., Cohen, J. D. & Botvinick, M. M. Dorsal anterior cingulate cortex and the value of control. Nat. Neurosci. 19, 1286–1291 (2016).

47. Kolling, N. et al. Value, search, persistence and model updating in anterior cingulate cortex. Nat. Neurosci. 19, 1280–1285 (2016).

48. Pochon, J.-B., Riis, J., Sanfey, A. G., Nystrom, L. E. & Cohen, J. D. Functional Imaging of Decision Conflict. J. Neurosci. 28, 3468–3473 (2008).

49. Botvinick, M. M., Cohen, J. D. & Carter, C. S. Conflict monitoring and anterior cingulate cortex: an update. Trends Cogn. Sci. 8, 539–546 (2004).

50. Niv, Y. Reinforcement learning in the brain. J. Math. Psychol. 53, 139–154 (2009).

51. O’Doherty, J. Dissociable Roles of Ventral and Dorsal Striatum in Instrumental Conditioning. Science 304, 452–454 (2004).

52. Marche, K., Martel, A.-C. & Apicella, P. Differences between Dorsal and Ventral Striatum in the Sensitivity of Tonically Active Neurons to Rewarding Events. Front. Syst. Neurosci. 11, (2017).

53. Morrison, I., Tipper, S. P., Fenton-Adams, W. L. & Bach, P. “Feeling” others’ painful actions: The sensorimotor integration of pain and action information. Hum. Brain Mapp. 34, 1982–1998 (2013).

54. Guo, X. et al. Empathic neural responses to others’ pain depend on monetary reward. Soc. Cogn. Affect. Neurosci. 7, 535–541 (2012).

55. Whitmarsh, S., Nieuwenhuis, I. L. C., Barendregt, H. P. & Jensen, O. Sensorimotor alpha activity is modulated in response to the observation of pain in others. Front. Hum. Neurosci. 5, (2011).

56. Jepma, M., Verdonschot, R. G., van Steenbergen, H., Rombouts, S. A. R. B. & Nieuwenhuis, S. Neural mechanisms underlying the induction and relief of perceptual curiosity. Front. Behav. Neurosci. 6, (2012).

57. Kruger, J. & Evans, M. The paradox of Alypius and the pursuit of unwanted information. J. Exp. Soc. Psychol. 45, 1173–1179 (2009).

58. Bromberg-Martin, E. S. & Hikosaka, O. Midbrain Dopamine Neurons Signal Preference for Advance Information about Upcoming Rewards. Neuron 63, 119–126 (2009).

59. Rodriguez Cabrero, J. A. M., Zhu, J.-Q. & Ludvig, E. A. Costly curiosity: People pay a price to resolve an uncertain gamble early. Behav. Processes 160, 20–25 (2019).

60. Oosterwijk, S. Choosing the negative: A behavioral demonstration of morbid curiosity. PLoS One 12, e0178399 (2017).

61. Hsee, C. K. & Ruan, B. The Pandora effect: The power and peril of curiosity. Psychol. Sci. 27, 659–666 (2016).

62. Noordewier, M. K. & van Dijk, E. Curiosity and time: from not knowing to almost knowing. Cogn. Emot. 31, 411–421 (2017).

63. Dickinson, A. & Balleine, B. The Role of Learning in the Operation of Motivational Systems. in Stevens’ Handbook of Experimental Psychology (eds. Pashler, H. & Gallistel, R.) (John Wiley & Sons, Inc., 2002). doi:10.1002/0471214426.pas0312

64. Litman, J., Hutchins, T. & Russon, R. Epistemic curiosity, feeling-of-knowing, and exploratory behaviour. Cogn. Emot. 19, 559–582 (2005).

65. Schonberg, T. et al. Selective impairment of prediction error signaling in human dorsolateral but not ventral striatum in Parkinson’s disease patients: evidence from a model-based fMRI study. Neuroimage 49, 772–781 (2010).

66. Takahashi, Y., Schoenbaum, G. & Niv, A. Silencing the critics: understanding the effects of cocaine sensitization on dorsolateral and ventral striatum in the context of an actor/critic model. Front. Neurosci. 2, 86–99 (2008).

67. van Lieshout, L. L. F., Vandenbroucke, A. R. E., Müller, N. C. J., Cools, R. & de Lange, F. P. Induction and Relief of Curiosity Elicit Parietal and Frontal Activity. J. Neurosci. 38, 2579–2588 (2018).

68. Di Domenico, S. I. & Ryan, R. M. The Emerging Neuroscience of Intrinsic Motivation: A New Frontier in Self-Determination Research. Front. Hum. Neurosci. 11, (2017).

69. Preuschoff, K., Quartz, S. R. & Bossaerts, P. Human Insula Activation Reflects Risk Prediction Errors As Well As Risk. J. Neurosci. 28, 2745–2752 (2008).

70. Alexander, W. H. & Brown, J. W. The Role of the Anterior Cingulate Cortex in Prediction Error and Signaling Surprise. Top. Cogn. Sci. 11, 119–135 (2019).

71. Silvia, P. J. What Is Interesting? Exploring the Appraisal Structure of Interest. Emotion 5, 89–102 (2005).

72. Silvia, P. J. Appraisal components and emotion traits: Examining the appraisal basis of trait curiosity. Cogn. Emot. 22, 94–113 (2008).

73. Noordewier, M. K. & van Dijk, E. Interest in Complex Novelty. Basic Appl. Soc. Psych. 38, 98–110 (2016).

74. Botvinick, M. M., Braver, T. S., Barch, D. M., Carter, C. S. & Cohen, J. D. Conflict monitoring and cognitive control. Psychol. Rev. 108, 624–652 (2001).

75. Shenhav, A., Botvinick, M. M. & Cohen, J. D. The Expected Value of Control: An Integrative Theory of Anterior Cingulate Cortex Function. Neuron 79, 217–240 (2013).

76. Murayama, K., FitzGibbon, L. & Sakaki, M. Process Account of Curiosity and Interest: A Reward-Learning Perspective. Educ. Psychol. Rev. 31, 875–895 (2019).

77. Zink, C. F., Pagnoni, G., Martin-Skurski, M. E., Chappelow, J. C. & Berns, G. S. Human Striatal Responses to Monetary Reward Depend On Saliency. Neuron 42, 509–517 (2004).

78. Krebs, R. M., Boehler, C. N., Roberts, K. C., Song, A. W. & Woldorff, M. G. The Involvement of the Dopaminergic Midbrain and Cortico-Striatal-Thalamic Circuits in the Integration of Reward Prospect and Attentional Task Demands. Cereb. Cortex 22, 607–615 (2012).

79. Ozono, H. et al. Magic Curiosity Arousing Tricks (MagicCATs): A novel stimulus collection to induce epistemic emotions. Preprint at PsyArXiv doi:10.31234/osf.io/qxdsn (2020).

80. Fastrich, G. M., Kerr, T., Castel, A. D. & Murayama, K. The role of interest in memory for trivia questions: An investigation with a large-scale database. Motiv. Sci. 4, 227–250 (2018).

81. Peirce, J. W. Generating stimuli for neuroscience using PsychoPy. Front. Neuroinform. 2, (2009).

82. Murayama, K., Matsumoto, M., Izuma, K. & Matsumoto, K. Neural basis of the undermining effect of monetary reward on intrinsic motivation. Proc. Natl. Acad. Sci. 107, 20911–20916 (2010).

83. Murayama, K. et al. How self-determined choice facilitates performance: A Key role of the ventromedial prefrontal cortex. Cereb. Cortex 25, 1241–1251 (2015).

84. Worsley, K. J. et al. A unified statistical approach for determining significant signals in images of cerebral activation. Hum. Brain Mapp. 4, 58–73 (1996).

85. Pauli, W. M., Nili, A. N. & Tyszka, J. M. A high-resolution probabilistic in vivo atlas of human subcortical brain nuclei. Sci. Data 5, 180063 (2018).

86. Mumford, J. A., Poline, J.-B. & Poldrack, R. A. Orthogonalization of Regressors in fMRI Models. PLoS One 10, e0126255 (2015).

87. Rissman, J., Gazzaley, A. & D’Esposito, M. Measuring functional connectivity during distinct stages of a cognitive task. Neuroimage 23, 752–763 (2004).

88. Göttlich, M., Beyer, F. & Krämer, U. M. BASCO: a toolbox for task-related functional connectivity. Front. Syst. Neurosci. 9, (2015).

89. Tomasi, D. & Volkow, N. D. Laterality Patterns of Brain Functional Connectivity: Gender Effects. Cereb. Cortex 22, 1455–1462 (2012).

90. Wang, D., Buckner, R. L. & Liu, H. Functional Specialization in the Human Brain Estimated By Intrinsic Hemispheric Interaction. J. Neurosci. 34, 12341–12352 (2014).

91. Barr, D. J., Levy, R., Scheepers, C. & Tily, H. J. Random effects structure for confirmatory hypothesis testing: Keep it maximal. J. Mem. Lang. 68, 255–278 (2013).

92. Brauer, M. & Curtin, J. J. Linear mixed-effects models and the analysis of nonindependent data: A unified framework to analyze categorical and continuous independent variables that vary within-subjects and/or within-items. Psychol. Methods 23, 389–411 (2018).

93. Murayama, K., Sakaki, M., Yan, V. X. & Smith, G. M. Type I error inflation in the traditional by-participant analysis to metamemory accuracy: A generalized mixed-effects model perspective. J. Exp. Psychol. Learn. Mem. Cogn. 40, 1287–1306 (2014).

94. Ludbrook, J. Interim analyses of data as they accumulate in laboratory experimentation. BMC Med. Res. Methodol. 3, 15 (2003).

95. Šidák, Z. Rectangular Confidence Regions for the Means of Multivariate Normal Distributions. J. Am. Stat. Assoc. 62, 626–633 (1967).

96. Stone, M. Comments on Model Selection Criteria of Akaike and Schwarz. J. R. Stat. Soc. Ser. B 41, 276–278 (1979).

97. Wagenmakers, E.-J. A practical solution to the pervasive problems of p values. Psychon. Bull. Rev. 14, 779–804 (2007).

98. Masson, M. E. J. A tutorial on a practical Bayesian alternative to null-hypothesis significance testing. Behav. Res. Methods 43, 679–690 (2011).

99. Bates, D., Mächler, M., Bolker, B. & Walker, S. Fitting Linear Mixed-Effects Models Using lme4. J. Stat. Softw. 67, 1–48 (2015).

100. Allen, M., Poggiali, D., Whitaker, K., Marshall, T. R. & Kievit, R. A. Raincloud plots: a multi-platform tool for robust data visualization. Wellcome Open Res. 4, 63 (2019).

101. Muthén, L. K. & Muthén, B. O. Mplus User’s Guide. Seventh Edition. (2012).

102. Enders, C. K. & Tofighi, D. Centering predictor variables in cross-sectional multilevel models: A new look at an old issue. Psychol. Methods 12, 121–138 (2007).

103. McNeish, D., Stapleton, L. M. & Silverman, R. D. On the unnecessary ubiquity of hierarchical linear modeling. Psychol. Methods 22, 114–140 (2017).

104. Arend, M. G. & Schäfer, T. Statistical power in two-level models: A tutorial based on Monte Carlo simulation. Psychol. Methods 24, 1–19 (2019).

105. Leong, Y. C., Hughes, B. L., Wang, Y. & Zaki, J. Neurocomputational mechanisms underlying motivated seeing. Nat. Hum. Behav. 3, 962–973 (2019).

106. Geuter, S., Qi, G., Welsh, R. C., Wager, T. D. & Lindquist, M. A. Effect Size and Power in fMRI Group Analysis. Preprint at bioRxiv doi:10.1101/295048 (2018).

