## Supplementary Information for "Shared striatal activity in decisions to satisfy curiosity and hunger at the risk of electric shocks"

### Supplementary Methods

**Electric Stimulation and Calibration: Determining Shock Threshold**
Electric shocks were delivered via an ADInstruments ML4856 PowerLab 26T Isolated Stimulator using an MLADDF30 stimulating bar electrode with 30mm spacing of 9mm contacts. During the calibration session, the bar electrode was attached to the participant's distal phalange of the ring finger of the non-dominant hand. Each participant's stimulation level was set by first exposing them to a stimulation of 2mA (15 Pulses at 20 Hz, with a pulse duration of 200 μs) and increasing the current in steps of 0.2mA, until a suitable participant-specific threshold was identified that was considered uncomfortable and unpleasant but not painful. Each time a shock was applied the participant was asked to report whether they thought the current shock was with less, more, or the same intensity as the previous one, and whether they would continue to the next level. The procedures stopped when the participant indicated an intensity to be 'unpleasant' at least twice. After a personalised threshold was identified, the participant was informed that the intensity of shock would vary slightly about the final level of shock they last experienced and that the amount (duration) of shock would depend on their performance on the actual task.

**Modified procedures for individual experiments**
*Modification for the follow-up behavioural experiment (on subjective outcome estimation):* This follow-up experiment aimed to evaluate if a person's perception of the probability of getting electric shock (or winning the gamble) is altered by the strong motivational force of curiosity or an external incentive.

In the experimental task, the display of a stimulus was followed by participants giving their rating of curiosity (to a magic trick) or desirability (to a food item) on each trial. A wheel of fortune (WoF) was then shown. This time, participants were first asked to indicate how likely they thought they would actually win/lose in that particular lottery. This was achieved by moving a cursor along a visual analogue scale with 'likely to win' and 'likely to lose' labelled at two extreme ends respectively. The cursor first rested midway between the two ends and participants moved it closer towards the end that was labelled 'likely to win' to indicate their perceived greater chance of winning over losing, and vice versa. After that, participants were asked to take a gamble to either accept or reject the lottery (given the same WoF). No outcome (of the lottery) was presented at the end of a trial in this follow-up experiment to prevent the potential bias that the presented outcomes might have caused on the participant's subjective probability.

*Modification for the 'magic version' fMRI experiment (ver 1):* After the calibration for electric stimulation and completing the practice trials, participants performed the main task inside the MRI scanner, during which their fingers were still attached with MRI-compatible stimulating electrodes.

Each experimental trial began with a fixation cross jittered with a 4±2 s duration and a letter cue for 1 s. Participants then (after a fixed interval of 0.5 s) saw either a video of a magic trick (variable durations) or an image of a food item (2.5 s). The stimulus presentation might be followed by a rating scale and (if this appeared) participants made a rating of curiosity about the presented magic trick, or a rating of desirability regarding the presented food item. Due to limited scanning time, this 3.5 s rating phase occurred in only 10% of all trials in the fMRI experiment, and its purpose was mainly to retain participants' attention to the stimuli during the task. Then, in the 4 s decision phase, a WoF was displayed and participants made a decision regarding whether to accept or reject the lottery. Before and after the decision phase, there was a blank screen temporarily jittered with a 2±0.5 s duration. At the end of each trial, the outcome of the participant's decision (either 'win' or 'loss' if accepting the gamble, or 'pass' if rejecting it) was shown for 2.5 s.

The task was split into two MRI scans, with equal numbers of curiosity and food trials in each scan (n=18 for each category in each scan). After completing the task in the MRI scanner, participants returned to the lab where they were shown all the stimuli again and required to rate each of them. For the magic tricks, participants also gave a subjective indication of the moment of surprise (i.e. the moment at which they thought the trick 'magically' unfolded, inducing surprise) while watching a video. Based on the responses from all participants, we obtained the group's mean moment of surprise for each trick and used it to indicate the onset of elicitation (of curiosity) in the fMRI experiment for each 'magic' curiosity trial.

*Modification for the 'trivia version' fMRI experiment (ver 2)*: The second version used a collection of trivia questions (instead of magic trick videos) as stimuli to induce curiosity. In terms of operational procedures, this experiment overall implemented similar procedures as the magic version. In each trial of the task, after seeing a fixation crosshair and a letter cue ('T' for trivia; 'F' for food), participants were shown either a trivia question or an image of food, for 3.5s. Again, there was a rating scale that occurred in only 10% of all trials. The rest of each trial, including the decision phase, the temporarily jittered blank screens before and after the decision phase, as well as the outcome presentation, were run in a similar manner to the first version.

Furthermore, with this second version, we adopted an online adjustment approach for the presentation of the WoF, with an aim to prevent participants from developing a tendency for certain decisions (i.e. either overly accepting or overly rejecting the gambles) across the task. More specifically, the task program computed the average acceptance rate (based on how often the participant chose to accept/reject the gambles) for every 10 stimuli presented for each category. These acceptance rates then determined the frequency of each of the five different variants of WoF that would be displayed within the next 10 trials of a category. If the acceptance rate was low (<30%), the program would select to present more 'high-win-low-loss' variants. That is, the '83.3%-win-16.7%-loss' and '66.7%-win-33.3%-loss' WoFs would be displayed three times more often than the 'low-win-high-loss' variants in the next 10 trials of a category. On the other hand, if the acceptance rate was high (>70%), there would be more 'low-win-high-loss' WoFs (the '16.7%-win-83.3%-loss' and '33.3-win-66.7%-loss' variants would be shown three times more than the 'high-win-low-loss' ones) in the next 10 trials of a category. These steps were repeated for every 20 trials (incorporating 10 stimuli from each category that were mixed and presented in a random order) until the end of the task. By ensuring that the participants would have made relatively even 'accept' and 'reject' decisions with this procedure, we aimed to maximise the statistical power of the fMRI analysis.

This experiment was also implemented in two MRI scans. After completing the fMRI task, participants were shown all the stimuli again and required to give a rating for each of them.

**fMRI preprocessing**

The SPM12 software (The Wellcome Trust Centre for Neuroimaging, London, UK; www.fil.ion.ucl.ac.uk/~spm) was used for preprocessing and statistical analyses of the imaging data. Data preprocessing began with spatial realignment of the echo-planar imaging (EPI) volumes to correct for movement artefacts [1] and this step estimated motion parameters of each volume. With this step, we also tried to identify scans that showed abrupt movements (> 3mm displacement between adjacent volumes in any motion direction). As a result, one participant from the 'trivia version' fMRI experiment (ver 2) was excluded before further data analysis. The motion correction was followed by co-registering the structural image (T1) to the EPI data and then segmenting the T1 image based on a multi-channel approach. The segmented grey and white matter images were used to generate a DARTEL template, to which the EPI images of all subjects were warped. Finally, the DARTEL template and the EPI images were normalised to Montreal Neurological Institute (MNI) standard space [2], re-sliced to 3 x 3 x 3 mm voxels, and smoothed using a 9 mm Gaussian kernel to account for residual inter-subject differences and to comply with the continuity assumption of random field theory [3].

The blood-oxygen-level dependent signals across scan runs (i.e. including both sessions) were concatenated and then modelled using the general linear model (GLM), implemented in SPM. Onsets of each event type were specified within the same regressor. In all our models, we included reaction time as a parametric modulator, an attempt to account for any potential effects of duration in responding. These regressors were then convolved with the canonical hemodynamic response function. We also entered motion parameters and scan sequence as regressors of no interest in all GLMs, as well as applied a high pass filter with a cut-off of 1/128Hz to remove low frequency fluctuations (typically associated with biological and scanner noise). The effects of each event type were estimated using a fixed-effects model. The output images from each subject were then entered into a random effects analysis.

#### Supplementary Tables

**Supplementary Table 1. Contributions of different predictors in modelling decision (i.e. choice to accept or reject the gamble) tested by generalised linear mixed-effects model (GLME)**. Shown in the table are the predictors' beta coefficients (i.e. fixed-effects coefficients) [square brackets denote 95% confidence interval], the corresponding exponentiated values of β, Z-values, and P-values. Analysis was performed separately for curiosity and food conditions in each experiment. The formulas reported here show the structure of random effects that were test, and we also performed sensitivity analyses to evaluate the reliability and robustness of the results. #Data from the two versions of the fMRI experiment were combined and the corresponding GLME controlled with an extra fixed-effect term for the effect of version.

| Predictors | Curiosity | | | | Food | | | |
|---|---|---|---|---|---|---|---|---|
| | **Estimate, β** | **exp(β)** | **Z-value** | **P-value** | **Estimate, β** | **exp(β)** | **Z-value** | **P-value** |
| **Initial Behavioural (N=32)** | Formula: Decision ~ rating + probability + rating*probability + (1 \| subj) + (-1 + rating \| subj) + (-1 + probability \| subj) | | | | | | | |
| - Rating | 1.158 [0.84 – 1.47] | 3.185 | 7.236 | <0.001 | 1.146 [0.85 – 1.45] | 3.145 | 7.493 | <0.001 |
| - Presented Shock Probability | -1.282 [-1.60 – -0.96] | 0.277 | -7.801 | <0.001 | -1.120 [-1.59 – -0.83] | 0.30 | -6.292 | <0.001 |
| - Rating*Probability | -0.078 [-0.18 – 0.02] | 0.925 | -1.536 | 0.125 | 0.059 [-0.02 – 0.14] | 1.06 | 1.441 | 0.150 |
| **Follow-up Behavioural (N=29)** | Formula: Decision ~ rating + probability + rating*probability + (1 \| subj) + (-1 + rating \| subj) + (-1 + probability \| subj) | | | | | | | |
| - Rating | 1.564 [1.26 – 1.87] | 4.777 | 9.958 | <0.001 | 1.782 [1.412 – 2.153] | 5.943 | 9.424 | <0.001 |
| - Presented Outcome Probability | -1.789 [-2.20 – -1.38] | 0.167 | -8.479 | <0.001 | -1.728 [-2.10 – -1.35] | 0.178 | -9.015 | <0.001 |
| - Rating*Probability | 0.010 [-0.14 – 0.16] | 1.010 | 0.136 | 0.892 | -0.119 [-0.26 – 0.02] | 0.888 | -1.702 | 0.089 |
| **fMRI experiments (N=61)** | #Formula: Decision ~ rating + probability + rating*probability + version + (1 \| subj) + (-1 + rating \| subj) + (-1 + probability \| subj) | | | | | | | |
| - Rating | 0.589 [0.49 – 0.69] | 1.801 | 11.328 | <0.001 | 1.136 [0.98 – 1.29] | 3.115 | 14.476 | <0.001 |
| - Presented Outcome Probability | -1.198 [-1.37 – -1.03] | 0.302 | -13.588 | <0.001 | -1.349 [-1.54 – -1.16] | 0.260 | -14.062 | <0.001 |
| - Rating*Probability | -0.022 [-0.07 – 0.03] | 0.978 | -0.847 | 0.397 | -0.100 [-0.16 – -0.04] | 0.905 | -3.390 | <0.001 |
| - Stimulus version (v1 vs v2) | -0.017 [-0.40 – 0.36] | 0.983 | -0.088 | 0.930 | -0.153 [-0.57 – 0.26] | 0.858 | -0.724 | 0.469 |

**Supplementary Table 2. At Elicitation phase (during stimulus presentation), the main effect of decision (Accepted > Rejected gambles).** Greater activation for accepted, relative to rejected, trials across incentive categories in both fMRI versions. The effect of version and category along with participants' characteristics (i.e. gender and subjective shock expectation) were accounted for in the GLM. Results of the ROI analysis in caudate nucleus, nucleus accumbens, and VTA/SN are shown at the top of the table, and results of the exploratory whole-brain analysis are shown at the bottom. The results presented here are thresholded at a statistical level of *P*<0.001 (uncorrected) with a cluster extent: k ≥ 5 voxels for ROI analysis k ≥ 50 voxels for whole-brain analysis. Shown in the first column are the cluster-level FWE-corrected P-values: * indicates clusters that survived an adjusted significance of *P*<0.0167 in ROI analysis or a significance of *P*<0.05 in whole-brain analysis. L – left; R – right.

**Region-of-interest Analysis**

| FWE-corr p-value (cluster-level) | Cluster Size | Peak z-score | MNI coordinate x | y | z | Brain regions |
|---|---|---|---|---|---|---|
| **Nucleus Accumbens** | | | | | | |
| 0.009* | 13 | 4.45 | -6 | 6 | -9 | L |
| 0.014* | 5 | 3.62 | 9 | 12 | -9 | R |

No clusters in caudate and VTA/SN survived the ROI analysis.

**Whole-brain Analysis**

| FWE-corr p-value (cluster-level) | Cluster Size | Peak z-score | MNI coordinate x | y | z | Brain regions |
|---|---|---|---|---|---|---|
| **Medial structures** | | | | | | |
| 0.156 | 73 | 4.45 | -6 | 6 | -9 | L Nucleus accumbens |
| | | 3.93 | 6 | 15 | -9 | R Nucleus accumbens |
| 0.243 | 57 | 4.16 | 27 | 0 | -27 | R Hippocampus and medial temporal lobe |
| **Parietal/Occipital cortex** | | | | | | |
| 0.152 | 74 | 3.9 | -15 | -54 | 15 | L Occipitoparietal fissure |

**Supplementary Table 3. At Elicitation phase, the interaction effects of Decision*Category.** The effect of experiment version and participants' characteristics (i.e. gender and subjective shock expectation) were accounted for in the model. Any results of the ROI analysis are shown at the top of the table, and results of the exploratory whole-brain analysis are shown at the bottom. The results presented here are thresholded at a statistical level of $P<0.001$ (uncorrected) with a cluster extent: $k \geq 5$ voxels for ROI analysis $k \geq 50$ voxels for whole-brain analysis. Shown in the first column are the cluster-level FWE-corrected $P$-values: * indicates clusters that survived an adjusted significance of $P<0.0167$ in ROI analysis or a significance of $P<0.05$ in whole-brain analysis. L – left; R – right.

***Interaction 1: (Food: Accept > Reject) > (Curiosity: Accept > Reject) or (Curiosity: Reject > Accept) > (Food: Reject > Accept)***

**Region-of-interest Analysis**

| FWE-corr p-value (cluster-level) | Cluster Size | Peak z-score | MNI coordinate x | y | z | | Brain regions |
|---|---|---|---|---|---|---|---|
| **Nucleus Accumbens** | | | | | | | |
| 0.014* | 5 | 3.67 | -6 | 9 | -9 | L | |

No clusters in caudate nucleus and VTA/SN survived the predefined thresholds.

**Whole-brain Analysis**

| FWE-corr p-value (cluster-level) | Cluster Size | Peak z-score | MNI coordinate x | y | z | | Brain regions |
|---|---|---|---|---|---|---|---|
| **Frontal cortex** | | | | | | | |
| 0.001* | 311 | 5.01 | -3 | 30 | -12 | | Bilateral medial orbital frontal cortex |
| 0.144 | 76 | 4.16 | -48 | -12 | 0 | L | insula |

***Interaction 2: (Curiosity: Accept > Reject) > (Food: Accept > Reject) or (Food: Reject > Accept) > (Curiosity: Reject > Accept)***

No clusters survived the set thresholds in the ROI and whole-brain analyses.

There are no significant effects observed regarding the interaction of Decision*Version at elicitation phase.

**Supplementary Table 4. At Elicitation phase (during stimulus presentation), the main effect of decision (Rejected > Accepted gambles).** Greater activation for rejected, relative to accepted, trials across incentive categories in both fMRI versions. The effect of version and category along with participants' characteristics (i.e. gender and subjective shock expectation) were accounted for in the GLM. Any results of the ROI analysis are shown at the top of the table, and results of the exploratory whole-brain analysis are shown at the bottom. The results presented here are thresholded at a statistical level of $P<0.001$ (uncorrected) with a cluster extent: $k \geq 5$ voxels for ROI analysis $k \geq 50$ voxels for whole-brain analysis. Shown in the first column are the cluster-level FWE-corrected *P*-values: * indicates clusters that survived a significance threshold of $P<0.05$ in whole-brain analysis. L – left; R – right.

**Region-of-interest Analysis**

No clusters survived the ROI analysis.

**Whole-brain Analysis**

| FWE-corr p-value (cluster-level) | Cluster Size | Peak z-score | MNI coordinate x | y | z | | Brain regions |
|---|---|---|---|---|---|---|---|
| **Frontal cortex** | | | | | | | |
| <0.001* | 585 | 5.06 | 9 | 39 | 57 | R | Dorsomedial prefrontal cortex |
| | | 4.32 | 27 | 51 | 36 | R | Lateral prefrontal cortex |
| 0.272 | 53 | 3.82 | 42 | 27 | 48 | R | Middle frontal area |
| **Parietal/Occipital cortex** | | | | | | | |
| 0.058 | 111 | 4.62 | 21 | -102 | 6 | R | Posterior occipital lobe |
| **Other regions** | | | | | | | |
| 0.003* | 247 | 4.23 | 33 | 21 | -12 | R | Insula |
| 0.19 | 66 | 3.97 | -27 | 24 | -6 | L | Insula |

**Supplementary Table 5. At Decision phase, the main effect of decision (Accepted > Rejected gambles).** Greater activation for accepted, relative to rejected, trials across incentive categories in both fMRI versions. The effect of version and category along with participants' characteristics (i.e. gender and subjective shock expectation) were accounted for in the GLM. Results of the ROI analysis in caudate nucleus, nucleus accumbens, and VTA/SN are shown at the top of the table, and results of the exploratory whole-brain analysis are shown at the bottom. The results presented here are thresholded at a statistical level of *P*<0.001 (uncorrected) with a cluster extent: k ≥ 5 voxels for ROI analysis k ≥ 50 voxels for whole-brain analysis. Shown in the first column are the cluster-level FWE-corrected *P*-values: * indicates clusters that survived an adjusted significance threshold of *P*<0.0167 in ROI analysis or a significance threshold of *P*<0.05 in whole-brain analysis. L – left; R – right.

**Region-of-interest Analysis**

| FWE-corrected P-value (cluster-level) | Cluster Size | Peak z-score | MNI coordinate x | y | z | Brain regions |
|---|---|---|---|---|---|---|
| **Caudate Nucleus** | | | | | | |
| <0.001* | 144 | 6.91 | 9 | 12 | 0 | R |
| 0.001* | 134 | 6.22 | -9 | 12 | -3 | L |
| **Nucleus Accumbens** | | | | | | |
| 0.005* | 23 | 6.19 | 9 | 12 | -6 | R |
| 0.004* | 24 | 5.92 | -9 | 12 | -6 | L |
| **VTA/SN** | | | | | | |
| 0.004* | 33 | 4.61 | 9 | -18 | -15 | R |
| 0.005* | 27 | 4.55 | -12 | -18 | -9 | L |

**Whole-brain Analysis**

| FWE-corrected p-value (cluster-level) | Cluster Size | Peak z-score | MNI coordinate x | y | z | Brain regions |
|---|---|---|---|---|---|---|
| **Medial structures** | | | | | | |
| <0.001* | 4850 | 6.91 | 9 | 12 | 0 | R Caudate nucleus |
| | | 6.28 | -9 | 6 | -3 | L Caudate nucleus<br>This cluster also extends into bilateral thalamus, VTA/SN, cingulate gyri, medial frontal cortex, right anterior insula, and right prefrontal lobe and inferior frontal gyrus |
| **Parietal/Occipital cortex** | | | | | | |
| <0.001* | 832 | 6.05 | -27 | -69 | -12 | L Posterior occipital |
| 0.005* | 177 | 5.14 | -6 | 6 | 72 | L Paracentral lobule |
| | | 4.34 | 6 | 6 | 69 | R Paracentral lobule |
| 0.041* | 103 | 4.29 | 15 | -63 | 30 | R Occipitoparietal fissure |

**Supplementary Table 6. At Decision phase, the main effect of decision (Rejected > Accepted gambles).** Greater activation for rejected, relative to accepted, trials across incentive categories in both fMRI versions. The effect of version and category along with participants' characteristics (i.e. gender and subjective shock expectation) were accounted for in the model. Any results of the ROI analysis are shown at the top of the table, and results of the exploratory whole-brain analysis are shown at the bottom. The results presented here are thresholded at a statistical level of $P<0.001$ (uncorrected) with a cluster extent: $k \geq 5$ voxels for ROI analysis $k \geq 50$ voxels for whole-brain analysis. Shown in the first column are the cluster-level FWE-corrected *P*-values. L – left; R – right.

**Region-of-interest Analysis**

No clusters survived the pre-defined thresholds.

**Whole-brain Analysis**

| FWE-corr p-value (cluster-level) | Cluster Size | Peak z-score | MNI coordinate x | y | z | Brain regions |
|---|---|---|---|---|---|---|
| **Occipital cortex** | | | | | | |
| 0.107 | 73 | 5.83 | 15 | -90 | 0 | R Lingual |

**Supplementary Table 7. At Decision phase, the interaction effects of Decision*Category.** The effect of experiment version and participants' characteristics (i.e. gender and subjective shock expectation) were accounted for in the model. Any results of the ROI analysis are shown at the top of the table, and results of the exploratory whole-brain analysis are shown at the bottom. The results presented here are thresholded at a statistical level of *P*<0.001 (uncorrected) with a cluster extent: k ≥ 5 voxels for ROI analysis k ≥ 50 voxels for whole-brain analysis. Shown in the first column are the cluster-level FWE-corrected *P*-values. L – left; R – right.

***Interaction 1: (Food: Accept > Reject) > (Curiosity: Accept > Reject) or (Curiosity: Reject > Accept) > (Food: Reject > Accept)***

No clusters survived the pre-defined thresholds in the ROI and whole-brain analyses.

***Interaction 2: (Curiosity: Accept > Reject) > (Food: Accept > Reject) or (Food: Reject > Accept) > (Curiosity: Reject > Accept)***

**Region-of-interest Analysis**

| FWE-corr p-value (cluster-level) | Cluster Size | Peak z-score | MNI coordinate x | y | z | Brain regions |
|---|---|---|---|---|---|---|

No clusters survived the pre-defined thresholds in the ROI analyses.

**Whole-brain Analysis**

| FWE-corr p-value (cluster-level) | Cluster Size | Peak z-score | MNI coordinate x | y | z | Brain regions |
|---|---|---|---|---|---|---|
| **Medial structures** | | | | | | |
| 0.238 | 50 | 4.85 | 12 | -27 | 24 | R Splenium |

**Supplementary Table 8. At Decision phase, the interaction effects of Decision*Version.** The effect of incentive category along with participants' characteristics (i.e. gender and subjective shock expectation) were accounted for in the model. Any results of the ROI analysis are shown at the top of the table, and results of the exploratory whole-brain analysis are shown at the bottom. The results presented are thresholded at a statistical level of $P<0.001$ (uncorrected) with a cluster extent: $k \geq 5$ voxels for ROI analysis $k \geq 50$ voxels for whole-brain analysis. Shown in the first column are the cluster-level FWE-corrected *P*-values: * indicates clusters that survived an adjusted significance of $P<0.0167$ in ROI analysis or a significance of $P<0.05$ in whole-brain analysis. L – left; R – right.

***Interaction 1: (fMRI 1: Accept > Reject) > (fMRI 2: Accept > Reject) or (fMRI 2: Reject > Accept) > (fMRI 1: Reject > Accept)***

**Region-of-interest Analysis**

| FWE-corr p-value (cluster-level) | Cluster Size | Peak z-score | MNI coordinate x | y | z | | Brain regions |
|---|---|---|---|---|---|---|---|
| **VTA & SN** | | | | | | | |
| 0.019 | 5 | 3.69 | -12 | -24 | -12 | L | |

No clusters in caudate and nucleus accumbens survived the pre-defined thresholds.

**Whole-brain Analysis**

| FWE-corr p-value (cluster-level) | Cluster Size | Peak z-score | MNI coordinate x | y | z | | Brain regions |
|---|---|---|---|---|---|---|---|
| **Medial structures** | | | | | | | |
| 0.036* | 107 | 4.83 | -18 | -27 | -15 | L | Hippocampus, extending upward to posterior cingulate |
| | | 3.66 | -12 | -30 | -3 | L | Thalamus |

***Interaction 2: (fMRI 2: Accept > Reject) > (fMRI 1: Accept > Reject) or (fMRI 1: Reject > Accept) > (fMRI 2: Reject > Accept)***

No clusters survived the pre-defined thresholds in the ROI and whole-brain analyses.

**Supplementary Table 9. Parametric modulation of activation by gamble decision at decision phase accounting for the presented outcome probability.** Greater activity in accepted (> rejected) trials across category in both fMRI versions. The effect of version and participants' characteristics (i.e. gender and subjective shock expectation) were accounted for. Results of the ROI analysis in caudate nucleus, nucleus accumbens, and VTA & SN are shown at the top of the table, and results of the exploratory whole-brain analysis are shown at the bottom. The results presented are thresholded at a statistical level of $P<0.001$ (uncorrected) with a cluster extent: $k \geq 5$ voxels for ROI analysis $k \geq 50$ voxels for whole-brain analysis. Shown in the first column are the cluster-level FWE-corrected *P*-values: * indicates clusters that survived an adjusted significance of $P<0.0167$ in ROI analysis or a significance of $P<0.05$ in whole-brain analysis. L – left; R – right.

**Region-of-interest Analysis**

| FWE-corr p-value (cluster-level) | Cluster Size | Peak z-score | MNI coordinate x | y | z | Brain regions |
|---|---|---|---|---|---|---|
| **Caudate structure** | | | | | | |
| 0.006* | 67 | 4.5 | 6 | 15 | 3 | R |
| 0.004* | 76 | 4.24 | -9 | 15 | 3 | L |
| **Nucleus Accumbens** | | | | | | |
| 0.013* | 7 | 3.81 | -6 | 6 | -6 | L |
| 0.013* | 7 | 3.7 | 9 | 9 | -6 | R |
| **VTA & SN** | | | | | | |
| 0.008* | 19 | 4.15 | 9 | -21 | -15 | R |
| 0.009* | 17 | 4 | -9 | -18 | -12 | L |

**Whole-brain Analysis**

| FWE-corr p-value (cluster-level) | Cluster Size | Peak z-score | MNI coordinate x | y | z | Brain regions |
|---|---|---|---|---|---|---|
| **Medial structures** | | | | | | |
| <0.001* | 603 | 4.59 | 9 | -27 | -18 | R VTA, also extending into left VTA |
| | | 4.5 | 6 | 15 | 3 | R Caudate |
| | | 4.24 | -9 | 15 | 3 | L Caudate, also extending into bilateral thalamus |
| 0.248 | 51 | 4.08 | 33 | -57 | -18 | R Medial inferior temporal gyrus |
| **Frontal cortex** | | | | | | |
| 0.062 | 95 | 5.13 | 48 | 3 | 51 | R Middle frontal (near premotor) area |
| 0.001* | 263 | 3.94 | 48 | 24 | 12 | R Inferior frontal lobe |
| **Parietal/Occipital cortex** | | | | | | |
| <0.001* | 889 | 6.65 | -18 | -84 | -9 | L Posterior occipital lobe |
| <0.001* | 296 | 4.64 | 27 | -93 | 9 | R Posterior occipital lobe |
| 0.104 | 78 | 4.54 | -6 | 6 | 72 | Bilateral paracentral lobule |

**Supplementary Table 10. Parameters and measures of paths in the multi-level mediation analysis.** The link between stimulus rating and choice (decision) were partially mediated by the activities of nucleus accumbens (NAcc) during the elicitation phase and caudate nucleus during the decision phase. The effects of the incentive category, experiment version, presented shock probability, as well as participants' characteristics (i.e. gender and subjective shock expectation) were accounted for in the model.

| | Estimate, β | Z-value | P-value |
|---|---|---|---|
| **Paths as parts of the proposed mediation model** | | | |
| Rating → NAcc (Elicitation Ph.) | 0.08 [0.03 – 0.12] | 3.38 | 0.001 |
| NAcc (Elicitation Ph.) → Caudate (Decision Ph.) | 0.03 [0.02 – 0.05] | 4 | <0.001 |
| Caudate (Decision Ph.) → Choice | 0.03 [0.01 – 0.04] | 3.50 | <0.001 |
| **Other direct paths** | | | |
| Rating → Choice | 0.34 [0.30 – 0.38] | 16.05 | <0.001 |
| NAcc (Elicitation Ph.) → Choice | 0.01 [-0.002 – 0.03] | 1.83 | 0.068 |
| Rating → Caudate (Decision Ph.) | 0.05 [0.01 – 0.08] | 2.53 | 0.011 |

**Supplementary Table 11. Brain regions showing decoupling effects with caudate nucleus in functional connectivity analysis.** Beta-series correlation with anatomical ROIs of left and right caudate revealed weaker connectivity in accepted compared with rejected gambles, at the decision phase. The effect of experiment version and incentive category along with participants' characteristics (i.e. gender and subjective shock expectation) were accounted for in the model. The results of the whole-brain analysis presented here are thresholded at a statistical threshold level of *P*<0.001 (uncorrected) with a cluster extent of k ≥ 20 voxels. Shown on the first column are the cluster-level FWE-corrected *P*-values: * indicates clusters that survived an adjusted significance of *P*<0.025 (due to multiple comparisons from different ROIs applied). L – left; R – right.

**Connectivity with Left Caudate for Accepted < Rejected gambles**

| FWE-corr p-value (cluster-level) | Cluster Size | Peak z-score | MNI coordinate x | y | z | | Brain regions |
|---|---|---|---|---|---|---|---|
| **Frontal-parietal region** | | | | | | | |
| 0.001* | 69 | 3.99 | -54 | 0 | 36 | L | Sensorimotor area (SMA) near inferior central sulcus and precentral gyrus |
| 0.159 | 23 | 4.37 | 42 | -18 | 30 | R | SMA near central sulcus and postcentral gyrus |
| **Occipital-parietal region** | | | | | | | |
| <0.001* | 192 | 4.15 | 21 | -54 | 48 | R | Cuneus and precuneus |
| <0.001* | 94 | 4.2 | -15 | -81 | 27 | L | Cuneus |
| 0.108 | 26 | 4.19 | -15 | -63 | 45 | L | Precuneus |

**Connectivity with Right Caudate for Accepted < Rejected gambles**

| FWE-corr p-value (cluster-level) | Cluster Size | Peak z-score | MNI coordinate x | y | z | | Brain regions |
|---|---|---|---|---|---|---|---|
| <0.001* | 79 | 4.48 | -36 | -15 | 30 | L | SMA inferior near central sulcus and postcentral gyrus |
| 0.022* | 40 | 4.07 | 6 | -42 | 39 | R | Posterior cingulate |
| 0.008* | 49 | 3.68 | 24 | -72 | 42 | R | Superior occipital area |

#### Supplementary References


1. Ashburner, J. & Friston, K. J. Rigid Body Registration. in *Human Brain Function* (eds. Frackowiak, R. S. J. et al.) 635–654 (Academic Press, 2003). doi:10.1016/B978-012264841-0/50034-2
2. Ashburner, J. & Friston, K. J. Spatial Normalisation Using Basis Functions. in *Human Brain Function* (eds. Frackowiak, R. S. J. et al.) 655–672 (Academic Press, 2003). doi:10.1016/B978-012264841-0/50035-4
3. Worsley, K. J. & Friston, K. J. Analysis of fMRI time-series revisited—Again. *Neuroimage* **2**, 173–181 (1995).
